# Complex I is essential for embryonic stem cell proliferation through NADH-mediated redox signaling

**DOI:** 10.64898/2026.10.26.761997

**Authors:** Tatiana Y. Starkova, Andrey A. Kuzmin, Igor B. Nazarov, Elena V. Skvortsova, Sergey V. Ponomartsev, Valentina Drianitchkina, Nikolay D. Aksenov, Alexey N. Tomilin, Sergey A. Sinenko

## Abstract

**Background:** Pluripotent embryonic stem cells (ESCs) possess unlimited self-renewal potential and the capacity to generate all cell types of an organism. Mitochondrial metabolism participates in the regulation of ESC self-renewal and differentiation, yet the specific roles of respiratory complex components in these processes remain largely unknown.

**Methods:** *Ndufs1*-knockout mouse ESCs were generated using the CRISPR/Cas9 method. Proliferation and cell cycle dynamics were assessed via flow cytometry with propidium iodide and Hoechst 33342/Pyronin Y staining. Mitochondrial physiology, including membrane potential, ROS levels, and mass was evaluated using JC-1, MitoSOX Red, DHE, and MitoTracker Red measurements. Cellular ATP and NAD/NADH levels were determined by luminescent and colorimetric assays. Pluripotency was confirmed by immunocytochemistry, Western blotting, teratoma formation, and directed differentiation into EpiSCs and cardiomyocytes. Downstream molecular and transcriptomic shifts were analyzed using RNA-seq and Western blotting.

**Results:** We demonstrate that while Complex I function is dispensable for the maintenance of pluripotent identity, its loss leads to a profound reduction of ESC proliferation rates. Deficiency of the core CI subunit Ndufs1 induces mitochondrial reductive stress, characterized by a decreased cellular NAD⁺/NADH ratio and elevated ROS levels. We find that cytosolic, but not mitochondrial, NADH accumulation acts as the primary signal limiting ESC proliferation, independent of energy depletion or excessive ROS. This process is associated with the activation of the integrated stress response and suppression of the NRF2/PGC-1α/SOD2 antioxidant axis.

**Conclusions:** Our findings identify NADH-driven reductive stress as a key checkpoint controlling ESC proliferation and reveal metabolic flexibility that preserves ESC developmental potential despite severe mitochondrial dysfunction. We propose that Complex I functions as a metabolic rheostat, tuning ESC mitotic activity through mitochondrial-to-nuclear redox signaling.

## Background

Embryonic stem cells (ESCs), derived from the epiblast, are defined as pluripotent due to their capacity to give rise to all types of cells of adult organism, but not extraembryonic tissues [1]. A hallmark of ESCs is their robust self-renewal ability, enabling indefinite proliferation in culture with retention of a multilineage differentiation capacity under defined conditions [2–5]. ESCs can be maintained in naive state or converted to primed pluripotency state, corresponding to pre- and post-implantation epiblast stages, respectively; the latter cells are referred to as epiblast stem cells (EpiSCs). Unlike naïve ESCs, primed EpiSCs exhibit a restricted differentiation capacity and are unable to contribute to mouse blastocyst chimeras. The transition of epiblast from naïve to primed pluripotency proceeds thorough a formative pluripotency state, represented *in vitro* by epiblast-like cells (EpiLCs). In contrast to primed EpiSCs, which rely predominantly on glycolytic metabolism, naïve ESCs utilize both glycolysis and oxidative phosphorylation (OXPHOS) for energy production [6–12]. The metabolic shift from a bivalent oxidative/glycolytic towards a primarily glycolytic metabolism is initiated early during the naïve-to-primed pluripotency transition, coincident with specification of EpiLCs [12]. Notably, activation of aerobic glycolysis due to stabilization of hypoxia-inducible factor-1α (HIF-1α) is sufficient to drive this transition [11]. Conversely, enhancing oxidative metabolism in primed EpiSCs can promote their reversal to a naïve-like pluripotent state [13]. Together, these findings underscore a critical mechanistic role for metabolic rewiring in regulating the transitions between distinct pluripotent states.

Beyond pluripotency state transitions, metabolic rewiring also governs another critical developmental process – embryonic diapause. This state of reversible developmental arrest requires strict metabolic control over cell proliferation at the blastocyst stage. During diapause, the epiblast exhibits significantly reduced proliferation while maintaining its pluripotent identity. Recent studies have highlighted the pivotal role of metabolic rewiring in this process, showing marked suppression of OXPHOS activity while preserving glycolysis and efficient nutrient utilization [14–16]. Specifically, inhibition of the Myc-controlled metabolic network or components of the L-leucine/mTOR signaling axis induces a diapause-like state in ESCs *in vitro* [16, 17]. However, whether direct suppression of the mitochondrial respiratory chain, particularly CI, is sufficient to trigger a diapause-like state in naïve ESCs remains an open question.

Cellular respiration occurs in mitochondria, with oxidative phosphorylation representing its final stage. This process is mediated by the electron transport chain (ETC) which comprises large transmembrane protein complexes embedded into the inner mitochondrial membrane (IMM). The sequential action of ETC complexes, which function as redox-driven proton pumps, couples oxygen consumption to adenosine triphosphate (ATP) synthesis. This electron flow is facilitated by four core complexes (Complexes I-IV) and mobile electron carriers, ubiquinone (coenzyme Q, CoQ) and cytochrome c (cyt c) [18, 19]. Complex I (CI), or NADH:ubiquinone oxidoreductase, is the central and most intricate multi-subunit complex of the mitochondrial electron transport chain. In mammals, CI is assembled from 45 subunits that are organized into two structural arms: a matrix arm, which protrudes into the mitochondrial matrix, and a membrane arm, embedded into the IMM spatially oriented approximately perpendicular to each other [20–25]. CI is functionally divided on three core modules: the N- and Q-modules mediating acceptance electrons from NADH and their transfer to ubiquinone (CoQ) [20, 21] and the P-module, or the membrane arm, acting as a proton pump, utilizing the energy derived from electron transfer to translocate protons across the IMM and thereby generate a proton gradient [26, 27].

The catalytic core of CI is composed of 14 highly conserved subunits. Seven core subunits of the matrix domain (Ndufs1, 2, 3, 7, 8, and Ndufv1, 2) are encoded by nuclear DNA. The seven core subunits (Nd1-6, and Nd4L) of the membrane domain are encoded by mitochondrial DNA (mtDNA) [28, 29]. The remaining 31 accessory subunits are not directly involved in the catalytic function of CI [30–33] but are essential for the assembly and structural stability of the holoenzyme [34, 35] and for modulating the rate of CI biogenesis [33]. Furthermore, certain accessory subunits have been shown to bind cofactors and participate in the regulation of cellular redox status [23, 24, 33, 36].

The function of each subunit is critical for complex activity, and the absence of any single core or accessory subunit results in embryonic lethality at stage E9.5 or within the first four weeks of life in mice [32, 37–41]. One established consequence of impaired CI function, achieved through inactivation of the core 75-kDa subunit (Ndufs1), is the generation of elevated levels of reactive oxygen species (ROS). In Drosophila, such ROS are directly involved in activation of the differentiation of hemocyte precursor cells, germline cells, and in regulation of the activity of hematopoietic niche cells controlling cellular immune response [42–44]. In mice, Ndufs1-mediated ROS generation contributes to the process of cellular reprogramming to a pluripotent state [45]. Furthermore, inactivation of the accessory subunit NDUFA7 in cardiomyocytes induces ROS generation that activates calcineurin signaling, leading to the expression of cardiac hypertrophy genes [35, 46]. Thus, through retrograde signaling mechanisms involving ROS and other secondary messengers, mitochondria regulate nuclear gene expression to direct diverse cell fates, including proliferation and differentiation [47–50].

However, the role of CI activity in regulating the biological processes of pluripotency maintenance and the exit from this state has remained unresolved. In this study, we investigated the function of the core CI subunit Ndufs1 in governing mouse ESC self-renewal and differentiation. Genetic inactivation of Ndufs1, and consequently, of the entire CI, resulted in a significant reduction in the proliferation of ESCs without compromising their pluripotent identity. Notably, our data suggest that CI deficiency might be involved in transitioning naïve ESCs into a metabolic state reminiscent of embryonic diapause.

## Methods

### Cell culture

Most cell culture media, supplements and other chemicals in this study were from <u>Thermo Fisher Scientific</u>, Waltham, MA, USA, unless otherwise specified. Mouse embryonic stem cells (ESCs, line E14 Tg2a, BayGenomics) were cultured on gelatin-coated plates in serum/LIF (SL) medium unless otherwise specified. The SL medium consisted of Knockout-DMEM supplemented with 15% fetal bovine serum (FBS, BioSera or HiClone, EU), 100 U/mL penicillin, 100 µg/mL streptomycin, 2 mM L-glutamine, 1× nonessential amino acids (NEAA), 50 µM β-mercaptoethanol (Sigma-Aldrich, USA), and in-house-produced human leukemia inhibitory factor (LIF) in dilution (1:5,000). Cells were passaged using a 0.05% Trypsin-EDTA. The medium was changed every other day, depending on cell density. To maintain the ground state of naïve pluripotency, ESCs were cultured on plates coated with 0.01% poly-L-ornithine in 2iL medium. This medium contained a standard N2B27 base (1:1 mix of Neurobasal and DMEM/F12), supplemented with 100 U/mL penicillin, 100 µg/mL streptomycin, 2 mM L-glutamine, 25 µM β-mercaptoethanol (Sigma-Aldrich, USA), as well as 1× N2, 1× B27, 3 µM CHIR-99021 (Axon Medchem, USA), 1 µM PD-0325901 (Axon Medchem, USA), and LIF (1:5,000). All cultures were maintained at 37°C in a humidified incubator with 5% CO₂. Human embryonic kidney cells (HEK-293T line) were grown in DMEM medium supplemented with 10% FBS (HyClone or BioSera, EU), 100 U/ml penicillin, 100 mg/ml streptomycin, 2 mM L-glutamine.

### Plasmid constructs for CRISPR/Cas9-mediated knockout of mouse Ndufs1

For CRISPR/Cas9-mediated knockout of the *Ndufs1* gene, we used a modified pX330-U6-Chimeric_BB-CBh-hSpCas9 plasmid, which encoded the Cas9 nuclease, a single-guide RNA (sgRNA) sequence, and enhanced green fluorescent protein (EGFP) for cell selection. sgRNA sequences were designed using the Benchling online platform (www.benchling.com). Two sgRNAs were used: 5’-gCCTATAAAAAGGGCCTTGAT-3’ (the ‘g’ was added at the 5’ end for efficient transcription from the U6 promoter), targeting the second exon; and 5’-gTGATGGCCAGTTCGCACAAC-3’ (with an added ‘g’ for the same purpose) (Fig. S1A). A scrambled control sequence (5’-GCACTACCAGAGCTAACTCA-3’) was also cloned. The sgRNA sequences were cloned into the pX330-U6-Chimeric_BB-CBh-hSpCas9 vector using BpiI restriction sites. Transient transfection of the pX330-U6-Chimeric_BB-CBh-hSpCas9_sgRNA vector was carried out in 24-well plates using FuGene HD transfection reagent (Promega, USA). Cells were seeded at a density of 1 × 10⁴ cells/cm² the day before transfection. One hour prior to transfection, the medium was replaced with serum-free OptiMEM supplemented with LIF. Twelve hours after transfection, the medium was changed to standard SL medium. On day 3 post-transfection, cells were detached using 0.05% Trypsin-EDTA, and the EGFP-positive population was sorted on a S3e cell sorter (Bio-Rad, USA). The sorted cells were seeded at low density (2,000 cells per 6-cm dish) onto gelatin-coated dishes and cultured under standard conditions until single colonies formed. Individual ESC colonies were then isolated and validated by Western blotting and genotyping.

### Genotyping of CRISPR/Cas9 knockout clones

To genotype CRISPR/Cas9 knockout ESC clones, primers were designed such that the 3’ end of one primer was complementary to the unmodified target sequence. This ensured that only the wild-type allele would be amplified, whereas DNA containing mutations in both Ndufs1 alleles would not produce a PCR product. Genotyping and subsequent analysis were performed for clones generated using sgRNA #1. The primer sequences were as follows: forward, 5’-AGGATACCTATAAAAAGGGCCTTGA-3’; reverse, 5’-ACAGTCATACAGCAGCGTCT-3’.

### Preparation lentiviruses mitoLbNOX or LbNOX

The LbNOX or mitoLbNOX sequences (Addgene # 75285 and 74448) [51] were cloned into the pLVTHM-IRES-Puro vector at the AscI/SpeI restriction sites. The empty pLVTHM-IRES-Puro was used as a control. A day before transfection, HEK293T cells were seeded in 10-cm dishes at a confluence of 30% and grown on DMEM-S medium containing DMEM, 10% fetal bovine serum (HyClone, EU), 100 U/ml penicillin, 100 mg/ml streptomycin, 2 mM L-glutamine in a CO_2_-incubator. The next day, 2 hours before transfection, medium was replaced with fresh medium without penicillin and streptomycin. Then, each vector of interest was co-transfected with pMD2 and psPax2 packaging vectors into Hek293T cells using 40 µg polyethylenimine (PEI) transfection reagent (PEI MAX® MW 40000, Polysciences, Inc., Warrington, PA, USA). After 18 hours the media was replaced with fresh DMEM-S and viral particles were collected from the media by centrifugation at 47 000g for 2h at 4°C [52]. Lentiviral pellets were resuspended in OptiMEM and stored at −80°C.

### Western blotting

Western blotting was performed as previously described [45]. Mouse ESCs were trypsinized, collected, and lysed in Laemmli loading buffer. Proteins were separated by 10% or 12% SDS-PAGE and transferred onto a nitrocellulose membrane (Amersham Hybond-ECL, USA) using a semi-dry transfer system (Helicon, Russia). Membranes were blocked in 3% skim milk in phosphate-buffered saline (PBS) containing 0.1% Tween-20 (PBST) for 1 h at room temperature and then incubated with primary antibodies (see Table 1) overnight at 4°C. Following three washes with PBST, membranes were incubated for 2 h with HRP-conjugated secondary antibodies (Jackson ImmunoResearch, USA, see Table 1) at room temperature (RT). After additional washing with PBST, protein bands were detected using West Dura Extended Duration Substrate and visualized on a ChemiDoc Touch Imaging System (Bio-Rad, USA). β-Tubulin or β-Actin were used as a loading control.

**Table 1.**
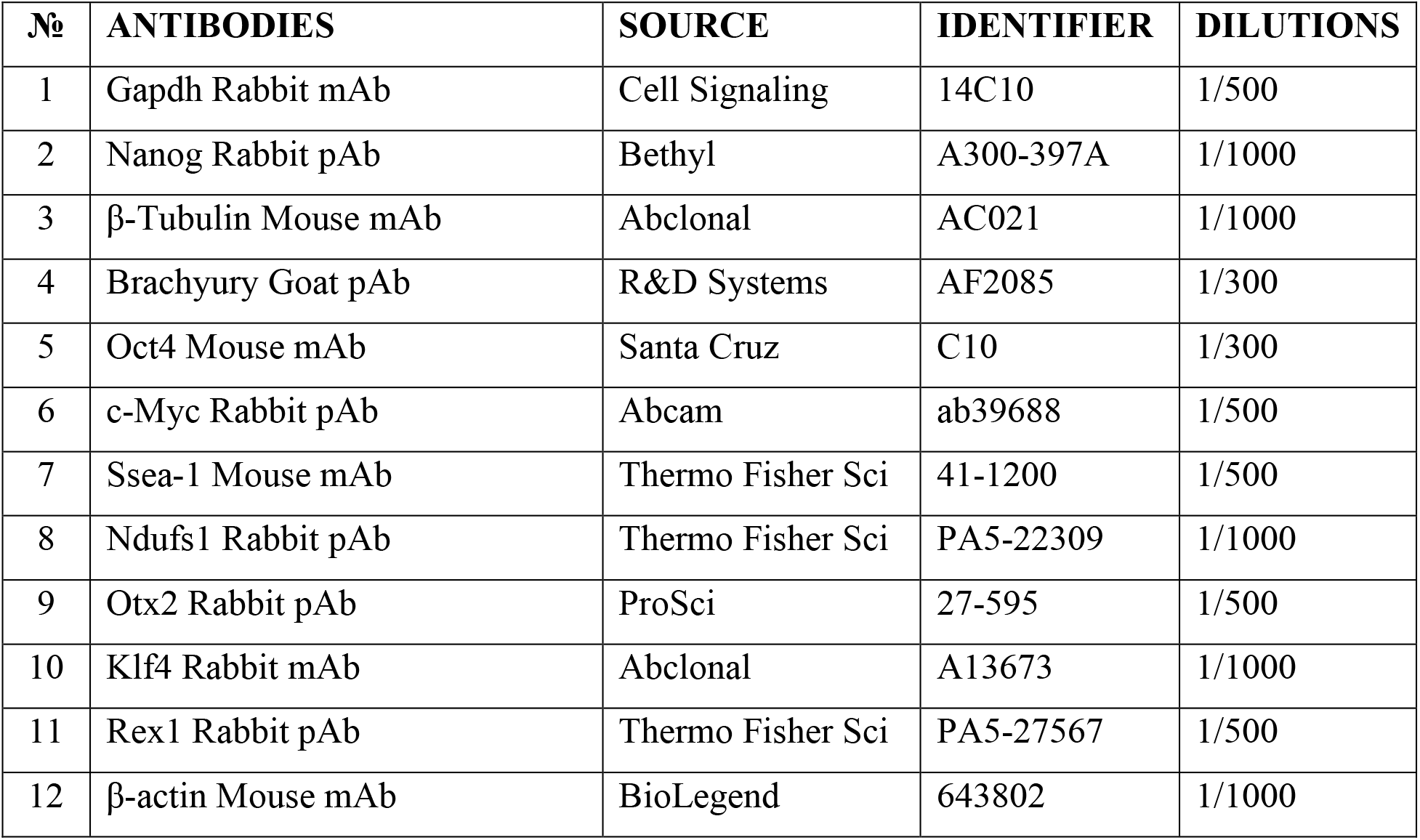
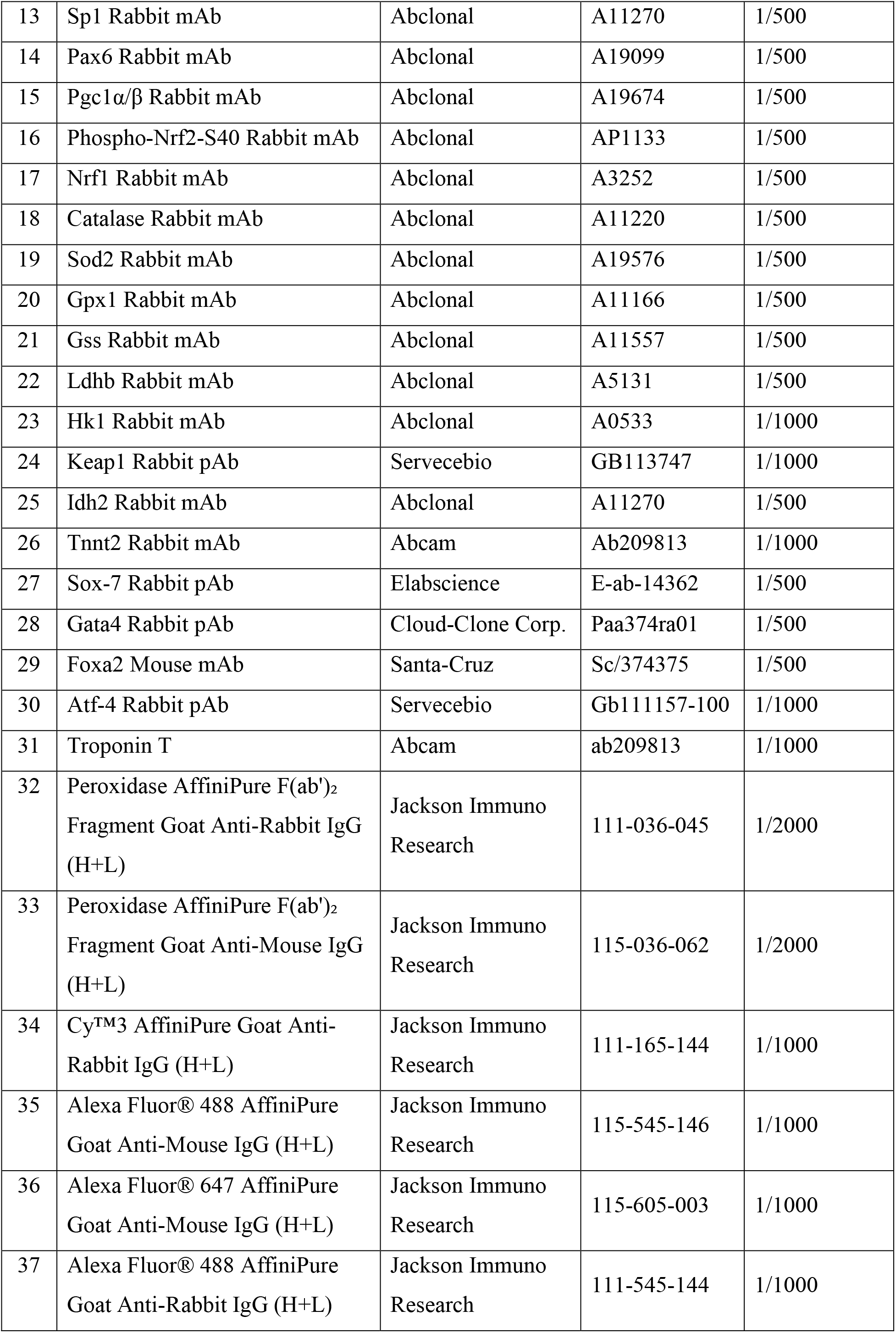
Antibodies.

## Antibodies

### Immunocytochemistry

Immunostaining was performed as previously described [53]. Cells were grown in 12-, 24-, or 48-well plates, fixed with 4% paraformaldehyde (PFA) in PBS for 10 min at RT, washed three times with PBS, and permeabilized with 0.1% Triton X-100 (AppliChem, USA) in PBS for 10 min. After blocking with 3% bovine serum albumin (BSA; VWR Chemicals, USA) in PBS for 1 h at room temperature, cells were incubated with primary antibodies (see Table 1) diluted in PBS containing 0.1% Tween-20 and 3% BSA overnight at 4°C. The following day, cells were washed three times with PBST and incubated with fluorescently labeled secondary antibodies (see Table 1) for 2 hrs at RT in the dark. After three washes with PBST, nuclei were counterstained with 1μM DAPI in PBS for 5 min at RT. Finally, cells were washed once with PBS and stored in PBS containing 0.1% sodium azide. Images were acquired using an EVOS cell imaging system (Thermo Fisher Scientific, USA).

### Assessment of cell death

To assess cell death, ESCs were collected 0.05% Trypsin-EDTA in PBS, mixed with 2-3 time-excess volume of SL medium, gently resuspended, centrifuged 3 min at 1200 rpm (ELMI CM-6M, Latvia), and cell pellet was collected (referred hereafter to as standard trypsinization). The cell pellet was then immediately resuspended in propidium iodide (PI, Invitrogen) solution (50 µg/mL in PBS). Following a 2-min incubation at RT in the dark, cells were washed with PBS and analyzed by flow cytometry on a CytoFLEX flow cytometer (Beckman Coulter, USA).

### Cell cycle analysis

Cells were seeded at a density of 1.5 × 10⁴ cells per well in a 12-well plate in SL medium. After 24 h, the medium was replaced with fresh medium containing 100 nM nocodazole (Sigma-Aldridge, #M1404, USA). Cells were cultured with nocodazole for 17 h, then washed and incubated in drug-free medium for an additional 24 h. To prepare cells for cell cycle analysis, they were washed with PBS, collected by standard trypsinization, and fixed in 100 µL of a staining solution containing RNase A (10 mg/mL), propidium iodide (PI, Sigma-Aldridge, #P4170, USA, 2 mg/mL), and 1% saponin in PBS. Cells were incubated in this solution for 40 min in the dark at RT. DNA content was measured by flow cytometry using a CytoFLEX flow cytometer (Beckman Coulter, USA), and the cell cycle phase distribution was determined using the analysis software.

### Hoechst 33342/Pyronin Y-based cell cycle analysis

To discriminate cells in the G0 resting phase of the cell cycle, the DNA and RNA content was measured by double staining using Hoechst 33342 (Thermo Fisher Scientific, #R37165, USA) and Pyronin Y (Sisco Research Laboratories, #75612, India) [54, 55]. Cells were seeded at a density of 1.5 x 10^4^ per well of a 12 well-plate in standard SL medium. After 48 hours, the cells were collected using a standard trypsinization protocol (in amount 4 x 10^4^ cells) and incubated with a staining solution containing 300 µl SL media and Hoechst 33342 (5 µg/ml) for 30 minutes in a CO_2_ incubator at 37°C in the dark, followed by an incubation with Pyronin Y (1 µg/ml) at 37°C for 10 min in the dark. Cells were then collected by centrifugation at 1200 x g for 3 min and resuspended in ice-cold PBS supplemented with 2% FBS and 25 mM HEPES (Thermo Fisher Scientific, #15630080, USA). Stained cells were analyzed within 30 minutes by a CytoFLEX flow cytometer (Beckman Coulter, USA).

### Animals and ethical statement

All animal procedures were approved by the Institutional Ethical Committee of the Institute of Cytology, Russian Academy of Sciences (Approval No. 07/25, approval date: Sept. 16, 2025; Project: “Investigation of the role of the Ndufs1 subunit of Complex I ETC in self-renewal and proliferation of mouse embryonic stem cells”) and complied with national legislation. Treatment-naïve female athymic BALB/c-Nude mice (*Crl:NU(NCr)-Foxn1nu*, The Jackson Laboratory, USA; 8–10 weeks old, 19–21 g; total N = 6) were housed under SPF conditions in groups of 6 in stainless steel cages (12-h light/dark cycle, 22 ± 2 °C, 45–65% humidity) with *ad libitum* access to food and water.

### Teratoma assay

An exploratory, biologically paired study design was used to minimize baseline variability and animal use in compliance with ethical guidelines for animal welfare. The sample size (N = 6) was chosen based on published teratoma experiments. Randomization for group allocation was not applicable since each mouse served as its own control. Animals were allocated to the study based on litter origin and similar body weight (±2 g), and co-housed to eliminate cage-location bias. Each mouse received bilateral subcutaneous injections into the hind limbs: 1 × 10⁶ control (Scr) ESCs into one side and 1 × 10⁶ *Ndufs1*-knockout (KO) ESCs into the contralateral side (total 12 injections, 6 per genotype). To account for clonal variation, three independent Scr clones and three KO clones were paired randomly; each pair was injected into two different mice as biological duplicates. Subcutaneous hind-limb injection was chosen as the standard, well-vascularized site allowing non-invasive monitoring.

The pre-established *a priori* inclusion criterion was successful teratoma formation (palpable tumor mass at the endpoint). Animal health, survival, and tumor burden were monitored daily using digital calipers. The experimental allocation was known only to the lead investigator during setup. Cell injections, harvesting, and histological processing were performed by a technician blinded to group identity. Blinding was lifted only after evaluation. At a predefined humane endpoint (when any tumor reached ∼1.5 cm; 3–4 weeks), mice were euthanized via rapid cervical dislocation by a trained technician without anesthesia. Teratomas were excised using aseptic surgical techniques. No animals or samples were excluded (n = 6 paired observations per genotype).

### Histological analysis of teratomas

Excised teratomas were fixed in 4% PFA overnight at 4 °C, dehydrated through a graded ethanol-isobutanol-xylene series, and embedded in paraffin. Five-micrometer sections were cut using a Leica RM2235 microtome, deparaffinized, rehydrated, and stained with hematoxylin and eosin according to standard protocols [56]. Slides were mounted in Canada balsam and analyzed using an EVOS Cell Imaging System microscope (Thermo Fisher Scientific, USA). Pluripotency was confirmed if at least three distinct representative tissue structures from each germ layer were identified per section. Because pluripotency was achieved in 100% of samples, this categorical parameter was not subjected to formal statistical analysis.

### RNA sequencing sample preparation

Scr and KO ESCs (passages 7–8) were seeded at a density of 1.8 × 10⁵ cells on 0.1%-gelatin-coated 6-cm dish. When cells reached ∼80% confluence, they were lysed directly in the dish with TRIzol Reagent and incubated for 5 min at RT. Lysates were collected, immediately frozen, and stored at –80°C. All samples were shipped on dry ice to Evrogen Joint Stock Company (Moscow, Russia) for subsequent RNA isolation, library preparation, and sequencing.

### RNA-seq data analysis and visualization

Differentially expressed genes (DEGs) were identified using a standard bioinformatics pipeline (e.g., DESeq2/edgeR). Volcano plots, bar graphs, and heatmaps were generated using exported z-scores or normalized read counts from the DEG analysis and plotted in GraphPad Prism. Pathway enrichment analysis was performed using the ReactomePA package in R, and results were visualized as ridge plots. Gene Ontology (GO) term semantic similarity analysis was conducted with REVIGO and visualized in R (v4.5.1).

### Flow cytometry analysis of superoxide levels and mass of active mitochondria

To assess mitochondrial superoxide and total cellular superoxide levels, and the mass of active mitochondria, cultured in 24-well plates cells in the logarithmic growth phase were washed with PBS, and incubated with either 5 µM MitoSOX Red (Thermo Fisher Scientific, USA), 1 µM dihydroethidium (DHE; Abcam, USA), or 100 nM MitoTracker red (Thermo Fisher Scientific, USA) in MEM medium for 30 min at 37°C in 5% CO₂ incubator. Following two washes with PBS, cells were collected by standard trypsinization, resuspended in PBS containing 2% FBS, and analyzed immediately on a CytoFLEX flow cytometer (Beckman Coulter, USA). Data were acquired and analyzed using CytExpert software.

### Measurement of ATP level in cultured ESCs

ATP levels were determined using the Luminescent ATP Detection Kit (ab113849, Abcam, USA). In brief, three control and three KO ESC cultures were seeded in 96-well plates at densities of 5 and 15×10^3^ cells per well and cultured under standard conditions for 48 h with a daily medium change. For the ATP assay, cells were lysed according to the manufacturer’s protocol for 5 min, and the substrate solution was added and incubated for 15 min at room temperature in the dark. In parallel, protein concentrations in the cell lysates were determined using the Pierce BCA Protein Assay Kit (Cat. No. 23225, Thermo Fisher Scientific, USA) according to the manufacturer’s instructions. Luminescence and absorbance were measured using a Varioskan LUX multimode reader (Thermo Fisher Scientific, USA) with three technical replicates per sample. ATP levels were normalized to the corresponding total protein content.

### Determination of the mitochondria membrane potential in ESCs

Mitochondrial membrane potential (MMP, ΔΨm) was determined using the JC-1 Mitochondrial Membrane Potential Assay Kit (Abcam, ab113850, UK). Three control and three KO ESC cultures were seeded in 12-well plates at densities of 1.5×10^4^ cells per well. The next day, cells were washed once with PBS, incubated with 300 µl of the 5 µM JC-1 working solution for 15 min in a CO_2_ incubator at 37°C, then washed twice with PBS and harvested by a standard trypsinization. Cells were resuspended in 150 µl PBS, and 1×10^4^ events per sample were analyzed by flow cytometry on a CytoFLEX equipment (Beckman Coulter, USA). Fluorescence was measured at an excitation wavelength of 475 nm and two emission filters for 530/30 (FITC channel, monomers) and 585/42 nm (PE channel, aggregates. MMP was assessed as the ratio of the median fluorescence intensities (MFI) in the PE and FITC channels (red/green ratio).

### Measurement of intracellular NAD levels

NAD and NADH levels were measured using the NAD/NADH Quantitation Colorimetric Kit (# MAK037-1KT, Sigma-Aldrich) according to the manufacturer’s instructions. Briefly, ESCs were collected by a standard trypsinization and counted. A pellet of 2.5 × 10⁵ cells was resuspended in 400 µL of NAD/NADH extraction buffer. Cells were homogenized by two freeze-thaw cycles (20 min on ice followed by 10 min at room temperature), vortexed for 10 s, and centrifuged at 10,000 × g for 10 min. To measure total NAD (NAD⁺ + NADH), 50 µL of the supernatant was mixed with 100 µL of NAD Cycling Enzyme Mix (98 µL Cycling Enzyme Buffer, 2 µL Enzyme Mix) in a well of a 96-well flat-bottom plate and incubated for 5 min at room temperature. Then, 10 µL of the colorimetric substrate was added to each well and incubated for 2 h at room temperature. For specific measurement of NADH, an aliquot of 200 µL of the supernatant was first incubated at 60 °C for 30 min to degrade NAD⁺ and then placed on ice for 2 min. This sample was centrifuged at 10,000 × g for 5 min. Next, 50 µL of the processed supernatant was added to 100 µL of NAD Cycling Enzyme Mix in a well of a 96-well plate, incubated for 5 min at room temperature, after which 10 µL of the substrate was added and incubated for 2 h at room temperature. The reaction was stopped by adding 10 µL of the fixative solution. Absorbance was read at 450 nm using a Thermo Scientific Varioskan LUX multimode reader (Thermo Fisher Scientific, USA). The total protein concentration in the lysates was determined using the Pierce™ BCA Protein Assay Kit according to the manufacturer’s protocol. The concentrations of total NAD and NADH were calculated from a standard curve and normalized to the total protein content (per mg of protein).

### Determination of the NAD+/NADH ratio in cells

The NAD/NADH-Glo™ Assay (Promega, USA) was used to determine the level of NAD/NADH in the samples. All manipulations with cells were carried out according to the manufacturer protocol. The protocol is designed to analyze cells in 50 µl of PBS per well in 96-well white luminometric plates. Each cell sample was divided into two: one sample was treated with acid to quantify NAD+ and the other was treated with base to quantify NADH. To a cell suspension of 10^4^ cells in 50 µl of PBS was added 0.2 N NaOH with 1% DTAB in a volume of 50 µl. To ensure cell homogeneity and lysis, the plate with cell lysates was subjected to short-term mixing on a shaker. After that, the entire volume of the lysate was divided into two samples. To the first (sample 1) was added 25 µl of 0.4 N HCl and heated for 15 minutes at 60°C. The second (sample 2) was also heated to 60°C for 15 minutes. Cool samples to RT, add 25 µl of 0.5 M Trizma base to each well of acid-treated cells (sample type 1) to neutralize the acid, and 50 µl of HCl/Trizma solution to each well containing base-treated samples (sample 2). The total sample volume per well of the plate is currently 100 µl. To perform the NAD/NADH-Glo™ assay, add an equal volume (100 µl) of NAD/NADH-Glo™ Detection Reagent directly to each well, mixing gently. After incubating the samples for 60 minutes, luminescence was recorded using a multimodal reader Thermo Scientific Varioskan LUX.

### Analysis of antioxidants and inhibitors on the ESC proliferation

To study the effect of low molecular weight inhibitors and antioxidants on proliferation, mouse ESCs were seeded at a density of 5×10^3^ per well in a 24-well plate and cultured for 48-72 hours in SL medium containing: 2 mM N-acetylcysteine (NAC, Sigma -Aldrich, USA) or 5 µM Ebselen (Ebs, Sigma-Aldrich, USA) or 25 µM MnTBAP (Abcam, UK) or 2.5 µM EUK134 (Abcam, UK) or 500-2000 nM ISRIB (ISR Inhibitor, InvivoChem, USA), 10 µM nicotinic acid (NA, Sigma-Aldrich, USA), or 10 mM dichloroacetate (DCA, Sigma-Aldrich, USA). After treatment, the cells were collected by standard trypsinization, washed once with PBS and after centrifugation the total cell pellet was resuspended in 100–200 µl of the serum free DMEM medium or PBS and analyzed by CytoFLEX flow cytometry equipment (Beckman Coulter, USA). Cell mortality was assessed by staining cells with 1 µg/ml propidium iodide (PI, Sigma-Aldrich, USA) for 1-2 min at RT. Cell death was monitored for each condition in proliferation assays, and there were no significant differences in cell mortality between treated and untreated cells unless mentioned otherwise.

### LbNOX and mitoLbNOX ectopic expression

ESCs were transduced by lentiviral particles encoding LbNOX, mitoLbNOX [51] or an empty control vector, and after 48 h cells were selected with puromycin (1 μg/ml). The puromycin-selected ESCs were seeded at a density of 5 x 10^3^ cells per well in a 24-well plate and cultured for 48-72 h in SL medium. Cells were harvested by standard trypsinization, washed once with PBS and collected by centrifugation. The total cell pellet was then resuspended in 100–200 µl of the serum free DMEM. To assess cell mortality, cells were stained with 1 µg/ml propidium iodide (PI, Sigma-Aldrich, USA) for 1-2 min at RT and immediately analyzed using a CytoFLEX flow cytometer (Beckman Coulter, USA).

### ESC differentiation into EpiSC cells

The week before differentiation, cells were cultured in the 2iL medium (see “Cell culture” section) on plates treated with 0.01% poly-L-ornithine (Sigma-Aldrich, USA) [56, 57]. On the first day of differentiation, cells were seeded at a density 100 x 10^3^ cells per well of 12-well plates pre-coated with fibronectin (17 µg/ml) in the differentiation medium containing 1:1 mixture of Neurobasal and DMEM/F12, supplemented with 1x N2, 1x B27 components, activin (0.02 µg/ml), bFGF (0.012 µg/ml), 0.01% KO-SR. Cells were cultured under these conditions for 10 days. On Day 4, 10 µM XAV was added to the medium. Cell cultures were re-seeded on day 3, 5 (for all lines), and 7 (for control lines only). Re-seeding (1:10 split ratio) was performed by treating the cells with collagenase-IV, followed by mechanical dissociation using a cell scraper. Differentiation progress was monitored by double immunocytochemical staining analysis using antibodies against Oct4 (Santa Cruz Biotech, USA) and Brachyury (AF2085, R&D Systems, USA), and the staining was visualized with an EVOS cell imaging system (Thermo Fisher Scientific, USA)

### Differentiation of ESCs into cardiomyocytes

Before differentiation, cells were cultured on SL medium (see “Cell culture” section) on a plate treated with 0.1% gelatin. Differentiation was carried out according to protocol, described elsewhere [58]. To initiate differentiation, cells were resuspended in the EB (embryonic body) medium DMEM, 15% FBS, 2 mM L-glutamine, 0.1 mM non-essential amino acids, 1 mM sodium pyruvate, 0.1 mM β-mercaptoethanol, 50 µg/ml L-ascorbic acid and cultured in 25-µl hanging drops on bacterial dish lids (1000 cells per drop) for 4 days. The bottom of these dishes was filled with PBS. After 4 days, embryonic bodies were transferred onto gelatin-treated 12-well tissue culture plates (7 per well) and cultured in the EB medium for another 6 days. The medium was changed daily. Differentiation efficiency was monitored by immunocytochemical analysis using antibodies against Troponin T (TnT, Abcam, UK), and the staining was visualized with an EVOS cell imaging system (Thermo Fisher Scientific, USA).

### Statistical analysis

All experiments were performed in at least three independent replicates. Data are presented as mean ± standard deviation (SD). Statistical significance between two groups was determined using the two-tailed unpaired Student’s t-test. A p-value < 0.05 was considered statistically significant.

## Results

### 1. *Ndufs1* deficiency impairs ESC proliferation

To investigate the role of ETC CI in self-renewal and differentiation of mouse ESCs, we inactivated *Ndufs1* gene using CRISPR/Cas9 technique. Three independent *Ndufs1* knockout clones (KO-3, -12, -18) and three scrambled control clones (Scr-1, -2, -3) were established (Fig. 1A). Frameshift mutations in the KO alleles were confirmed by DNA sequencing (Fig. S1A). Both KO and Scr ESC lines maintained normal karyotype (Fig. S1B). The cell and colony morphology of the generated KO ESC lines was indistinguishable from that of control Scr ESCs (Fig. S1C).

**Fig. 1.**
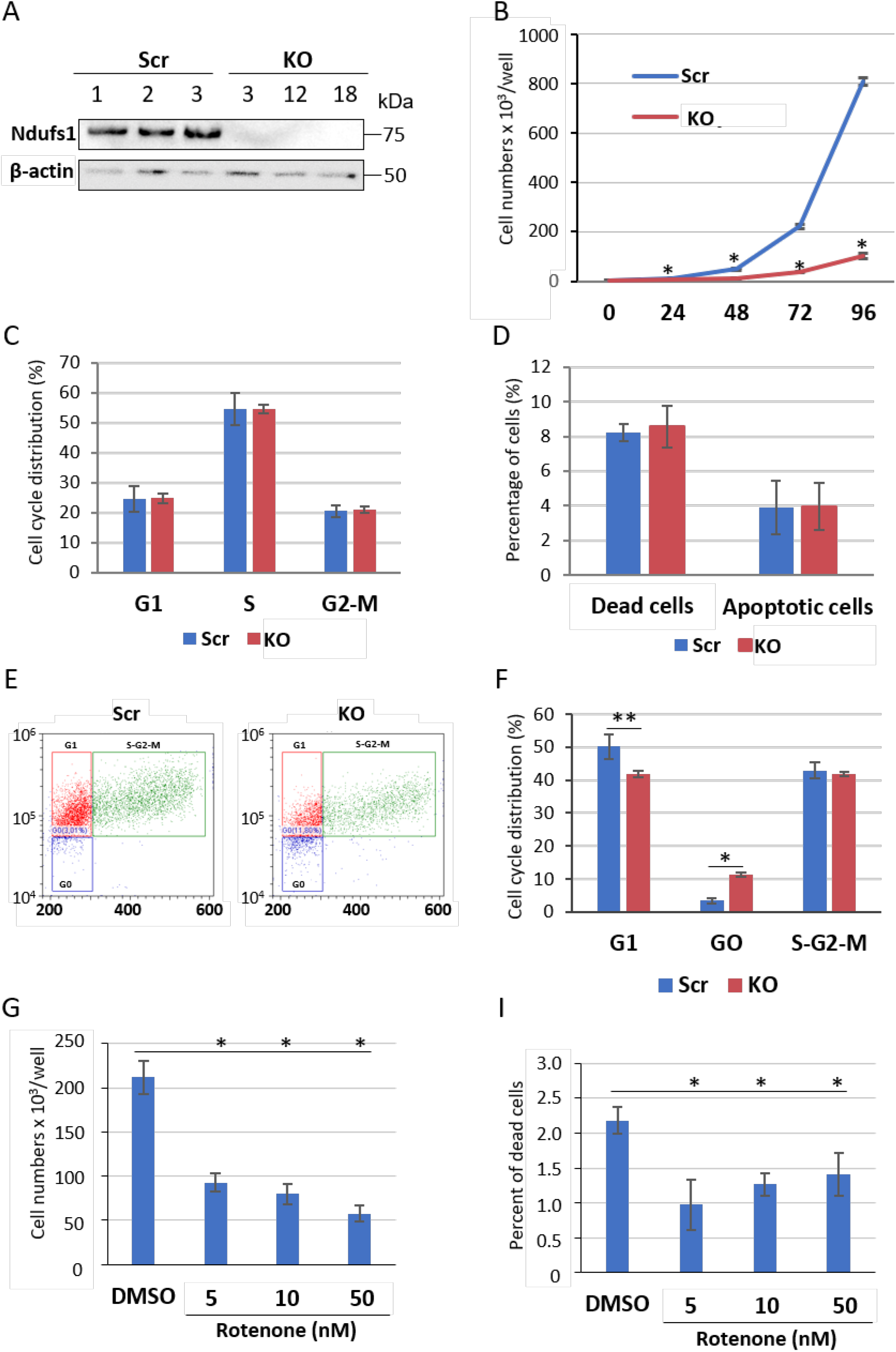
Genetic inactivation of *Ndufs1* significantly suppresses mouse ESC proliferation without compromising viability and the cell cycle progression. (A) Western blot analysis of Ndufs1 protein expression in control (Scr) and *Ndufs1* knockout (KO) ESC clones. β-Actin was used as a loading control. (B) Growth curves of Scr and KO ESCs over 96 hours. KO ESCs exhibited a 5- to 8-fold reduction in proliferation compared to Scr ESCs; data are presented as mean +/- SD; n=3 (biological replicates); *p<0.001. (C) Flow cytometry analysis showing unaltered cell cycle phase distribution in KO ESCs compared to Scr controls; mean +/- SD; n=3. (D) Quantification of dead and apoptotic cells using propidium iodide (PI) and Annexin V staining, respectively, followed by flow cytometry analysis. The proportions of dead or apoptotic cells remained unchanged in KO vs Scr ESCs; mean +/-SD; n=3. (E, F) Flow cytometry Hoechst 33342/Pyronin Y-based analysis of the G0 phase: representative plots (E) and quantifications (F), showing a significant increase in the proportion of resting cells in the G0 phase (blue gates in E) in KO ESCs compared to Scr controls under 2iL conditions; mean +/- SD; n=3 (G) Proliferation assessment following pharmacological CI inhibition. Treatment with sublethal concentrations of rotenone for 72 hours significantly reduced ESC numbers compared to the vehicle (DMSO)-treated control; means +/-SD; n=3 (biological replicates); *p<0.001. (I) Viability assessment under the same treatment as in G. The percentage of dead (PI-positive) cells was significantly reduced across all tested rotenone concentrations; mean +/-SD; n=3 (biological replicates); *p<0.001.

To evaluate the proliferative capacity of KO ESCs, cells were cultured under standard serum-based (serum/LIF, SL) conditions for 96 hours, and cell numbers were quantified at 24-hour intervals. Growth curve analysis revealed that KO ESCs proliferated 5- to 8-fold more slowly than control ESCs (Fig. 1B). During the first 24 hours, KO ESCs exhibited minimal proliferation, with cell numbers increasing by only 8.3%, compared to a 136.6% increase in control ESCs. Notably, the impaired proliferation of KO ESCs was not accompanied by alterations in cell cycle phase distribution when grown under SL culture conditions (Fig. 1C). Furthermore, *Ndufs1* inactivation did not increase cell death, including apoptosis (Fig. 1D). However, we observed a significant alteration in the cell cycle progression of KO ESCs when grown in serum-free (2iL) ground state conditions. Specifically, in comparison with control Scr ESCs, KO ESCs showed a significant (3-fold) increase in the proportion of cells residing in the G0 phase and a decrease in cells residing in G1 phase (∼16%) as revealed by Pyronin Y/ Hoechst 33342 analysis (Fig. 1 E, F). This suggests that the KO ESCs under the synchronized 2iL conditions tend to remain in a quiescent state, being arrested in the G0 phase, as previously shown for ground-state (rather than naïve-state) quiescent ESCs (2iL conditions) with inactivated histone acetyltransferase MOF [55]. Collectively, these results demonstrate that the genetic inactivation of *Ndufs1* under SL culture conditions slows proliferation without compromising cell viability.

Consistent with the above findings, pharmacological inhibition of CI function with 5, 10, and 50 nM rotenone also significantly reduced ESC proliferation, resulting in a 2- to 3.5-fold reduction in cell number after 72 hours compared to vehicle-treated (DMSO) control (Fig. 1G). In contrast to Ndufs1 inactivation, rotenone treatment reduced cell death rates (Fig. 1I), a discrepancy that warrants further investigation. These results suggest that ablation of CI function in ESCs induces a state of mitotic slowdown.

### 2. Ndufs1 is dispensable for ESC differentiation

To assess the pluripotency status of KO ESCs under SL conditions, the expression of key pluripotency markers was analyzed by Western blotting and immunocytochemistry. The analyses revealed that Oct4, Nanog, Rex1, and Ssea1 proteins levels were comparable between KO and Scr ESCs (Fig. 2A, B and Fig. S1C, D). RNA-seq analysis further confirmed that the transcript levels of these pluripotency factors were unchanged in KO ESCs (Supplementary Dataset 1). Klf4 represented a notable exception: its protein levels increased approximately two-fold (Fig. 2A, B), despite no change at the transcript levels (Supplementary Dataset 1), suggesting post-transcriptional regulation. Furthermore, transcriptome profiling revealed significant downregulation of genes associated with primed pluripotency in KO ESCs (Fig. S2A), indicating that CI deficiency reinforces a naïve pluripotent identity. To evaluate *in vivo* differentiation potential, we performed teratoma formation analysis. Both KO and control ESC clones successfully formed teratomas in immunodeficient nude mice, and histological analysis confirmed the presence of tissues representing all three germ layers in both KO and Scr ESC-derived teratomas (Fig. 2C). While no significant differences in teratoma size or weight were observed, KO ESC-derived teratomas exhibited a distinct morphology characterized by hollow, translucent, fluid-filled structures with extensive cavities surrounded by thin tissue layers (Fig. S2E). This altered morphology suggests that some perturbation in the CI-deficient cells may impair tissue organization and/or function. Such alterations may involve cell non-autonomous mechanism associated with CI deficiency, as previously reported [42, 44, 59], which warrants further investigation.

**Fig. 2.**
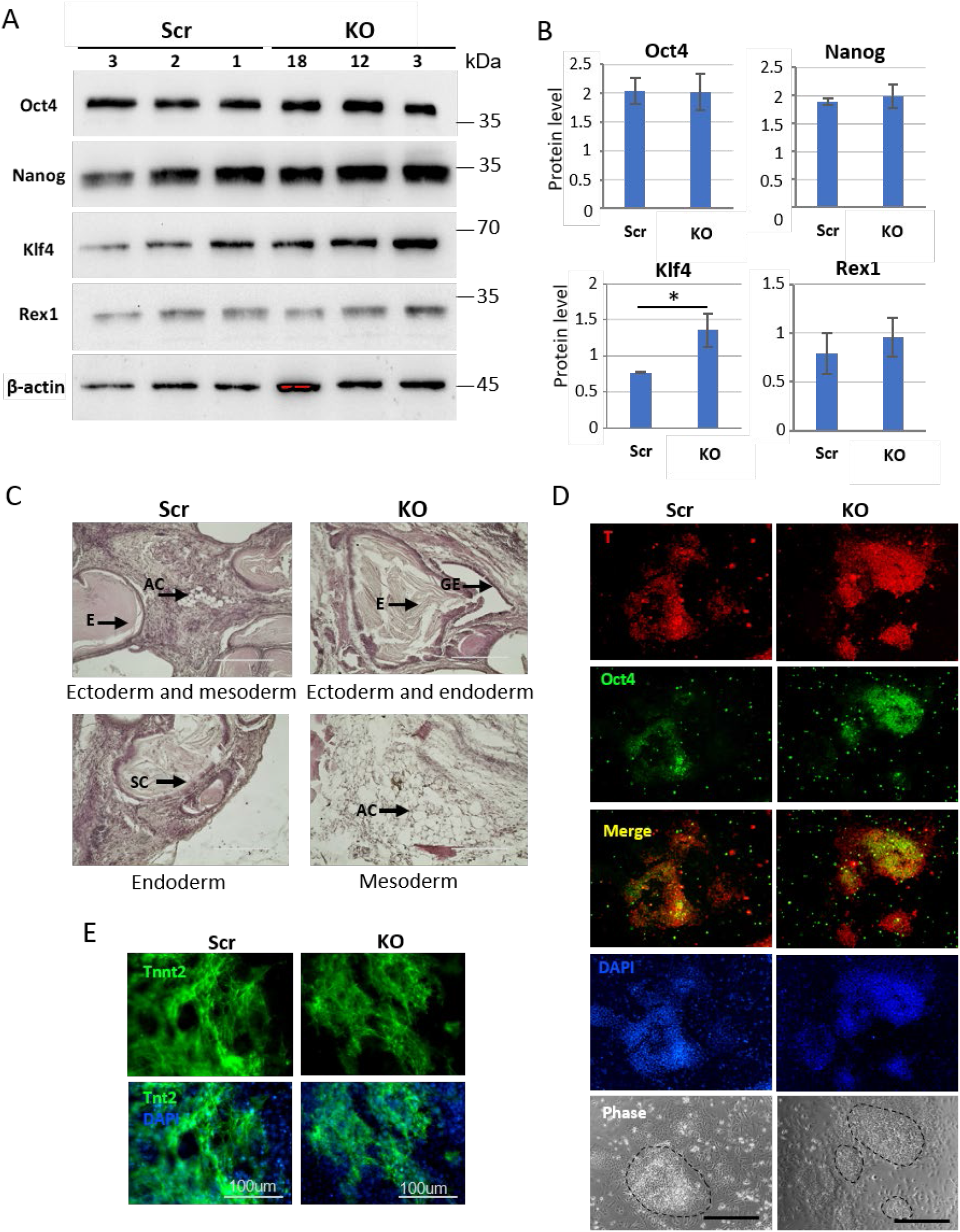
Ndufs1-deficient ESCs exhibit mostly unaltered expression of pluripotency markers and differentiation capacity. (A) Western blot analysis of core pluripotency factors in three independent control (Scr) and three KO ESC clones. Blots were probed with antibodies against Oct4, Nanog, Klf4, and Rex1. β-actin was used as a loading control. (B) Quantification of the protein marker expression in A, normalized to β-actin; mean ± SD; n=3 independent clones, *p<0.001. (C) KO ESCs retain the capacity to differentiate into tissues of all three germ layers. Histological analysis of teratomas derived from Scr and KO ESCs revealed representative tissues: adipocytes (ad, mesoderm), epidermis (ep, ectoderm), and intestinal epithelium and secretory cells (ie and sc, endoderm) (indicated by arrows). Scale bar: 200 μm. (D) KO ESCs differentiated into EpiSCs with efficiency comparable to Scr controls. Immunocytochemistry for the markers Brachyury (T, red) and Oct4 (green) confirms successful differentiation (Day 10) into EpiSCs of both KO and Scr ESCs. Nuclei were counterstained with DAPI (blue). Scale bar: 400 µm. (E) KO ESCs differentiated into cardiomyocytes with efficiency comparable to Scr controls. Immunofluorescence analysis for the cardiomyocyte marker cardiac Troponin T (Tnt2, green) confirms successful differentiation in both KO and Scr cells at Day 9. Nuclei are counterstained with DAPI (blue). Scale bar: 100 µm.

We next investigated the capacity of naïve KO ESCs to convert into epiblast stem cells (EpiSCs) *in vitro*, following an established protocol [60–64]. This naïve-to-primed pluripotency transition occurs over 10 days and proceeds through a transient formative pluripotency state (Day 2) represented by epiblast-like cells (EpiLCs). By Day 2, KO EpiLCs had reached a density comparable to that of controls. However, consistent with the reduced proliferation rates of KO ESCs, KO cultures exhibited significantly lower cell number from Day 3 to Day 7 (Fig. S2F). Despite this proliferative slowdown, KO cells successfully differentiated into EpiSCs by Day 10, as confirmed by the co-expression of Oct4 with the primed-state marker Brachyury (T) (Fig. 2D). These results demonstrate that while KO cells exhibit markedly reduced proliferation during the naïve-to-primed transition, they retain the capacity to reach the primed pluripotency state.

We next evaluated the ability of KO ESCs to differentiate into mature cardiomyocytes *in vitro* using an established protocol [65, 66]. Despite the reduced proliferation, KO ESCs successfully generated cardiomyocytes with kinetics comparable to that of control cells. By Day 9 of differentiation, both Scr- and KO-derived cardiomyocytes exhibited robust expression of the mature cardiac marker troponin T (Fig. 2E, Fig. S2G), indicating successful terminal differentiation. These results demonstrate that Ndufs1 inactivation slows proliferation but does not impair the potential of mouse ESCs to differentiate into either primed pluripotent stem cells or terminally differentiated cardiomyocytes *in vitro*. Collectively, these findings indicate that Ndufs1 function is dispensable for the execution of major differentiation programs in mouse ESCs.

### 3. *Ndufs1* loss induces metabolic stress in ESCs

We next analyzed key metabolic parameters of KO ESCs. As anticipated, mitochondrial membrane potential (MMP, ΔΨm) was markedly reduced, exhibiting an approximately fourfold reduction compared to control cells (Fig. 3A). MitoTracker Red (MTR) staining revealed ∼16.5% decrease in mitochondrial mass in KO ESCs relative to control cells (Fig. 3B). Despite the loss of CI function, total cellular ATP levels increased by ∼8% in KO ESCs compared to Scr controls (Fig. 3C), suggesting compensatory upregulation of glycolytic or alternative ATP-producing pathways in response to CI deficiency.

**Fig. 3.**
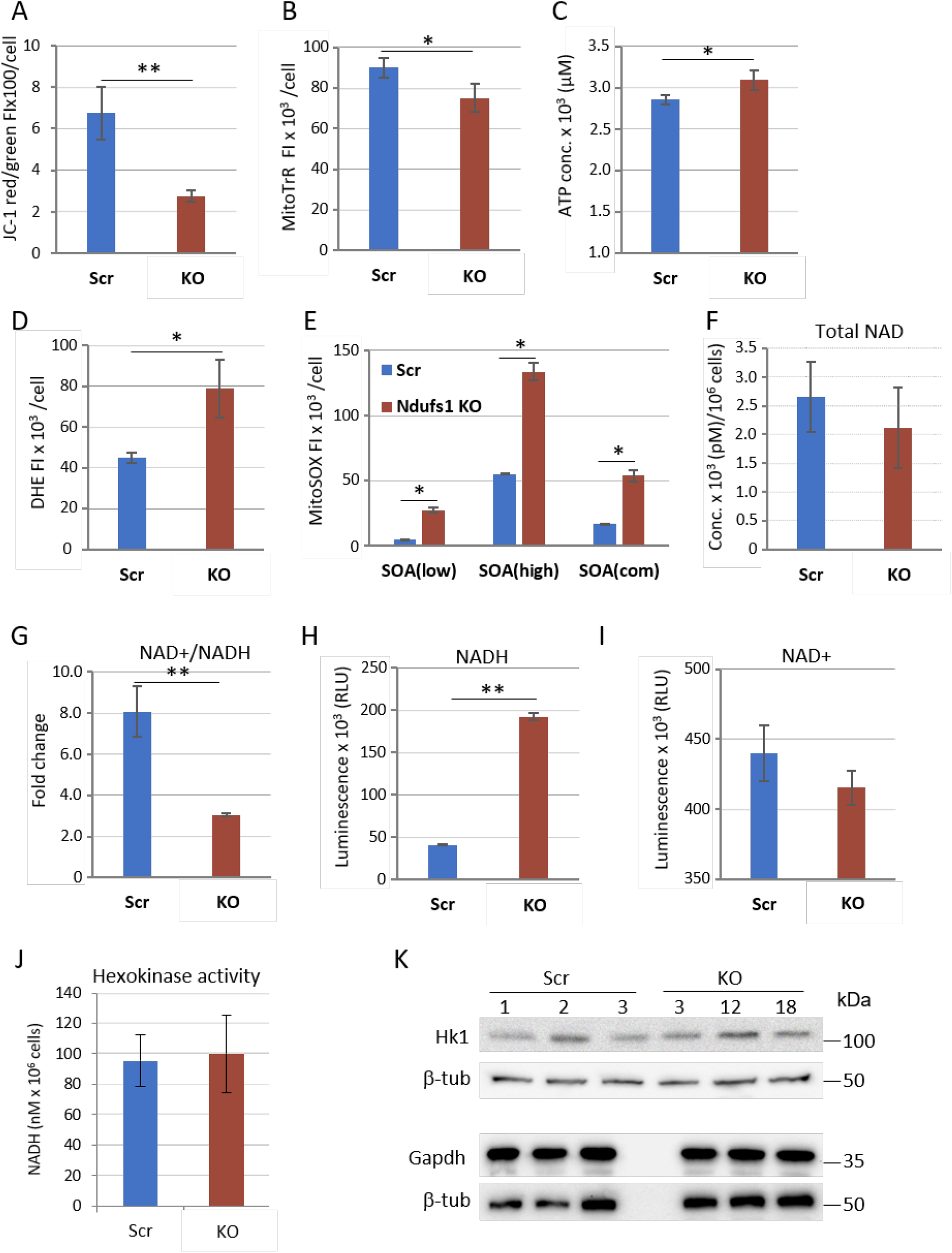
Ndufs1 deficiency induces mitochondrial depolarization coupled with an altered NAD+/NADH redox state and oxidative stress in ESCs. A) Mitochondrial membrane potential (MMP) was reduced in KO ESCs, compared to Scr controls, as measured by JC-1 staining and flow cytometry; mean fluorescence intensity (FI) per cell ± SD; n=3 (independent clones), **p<0.005. B) Mitochondrial mass was decreased in KO ESCs, compared to Scr controls, as assessed by MitoTracker Red (MTR) staining and flow cytometry; mean fluorescence intensity (FI) per cell ± SD; n=3 (independent clones), *p<0.05. C) Total cellular ATP levels were increased in KO ESCs relative to Scr controls; mean ± SD; n=3 independent clones, *p<0.05. D) Total cellular ROS levels were increased in KO ESCs, as measured by DHE staining and flow cytometry; mean fluorescence intensity per cell ± SD; n=3 (independent clones), *p<0.01. E) Mitochondrial ROS level elevation in KO ESCs, as revealed by MSR staining and flow cytometry; mean fluorescence levels per cell ± SD; n=3 (independent clones), *p<0.01. F) Unchanged total NAD levels in KO ESCs vs Scr ESCs; mean ± SD; n=6. G) The NAD+/NADH ratio was significantly reduced in KO ESCs relative to Scr controls; mean ± SD; n=6, **p<0.001. H) Strong elevation of NADH levels in KO ESCs, compared to Scr controls; mean ± SD; n=6, p*<0.001. I) Non-significant change of NAD+ levels in KO ESCs; mean ± SD; n=6. J) Unchanged glycolytic activity, as evidenced by total hexokinase activity, in KO ESCs; mean ± SD; n=6. K) The expression of the glycolytic enzymes hexokinase-1 (Hk-1) and glyceraldehyde-3-phosphate dehydrogenase (Gapdh) was unaltered in KO ESCs (n=3 independent clones), as shown by Western blot analysis. β-Tubulin (β-tub) was used as a loading control.

As expected, Ndufs1 inactivation in ESCs resulted in enhanced generation of mitochondrial and total cellular superoxide anion (superoxide, O₂•⁻), as evidenced by elevated fluorescence intensity of both dihydroethidium (DHE; ∼2-fold increase) and MitoSOX Red (MSR; ∼3-fold increase) in KO ESCs relative to control cells (Fig. 3D, E and Fig. S3A,B). Interestingly, MSR staining identified two distinct cellular subpopulations in both control and KO ESC clones, characterized by low and high superoxide levels (Fig. 3E and Fig. S3C). Superoxide production was significantly elevated in both subpopulations of KO ESCs (Fig. S3C), indicating that inactivation of the core CI subunit Ndufs1 leads to a substantial increase in ROS production.

CI dysfunction typically results in NADH accumulation due to loss of NADH-dehydrogenase activity [67, 68]. We have revealed that the total NAD pool was largely unchanged in KO ESCs compared to control cells (Fig. 3F). However, the NAD^+^/NADH ratio was reduced approximately fivefold in KO ESCs (Fig. 3G), primarily due to a substantial increase in NADH levels (Fig. 3H). In contrast, NAD^+^ levels were only marginally reduced in KO ESCs (Fig. 3I). These data indicate that NADH accumulation of in KO cells results from impaired CI activity, inducing a state of reductive stress within the mitochondrial matrix.

To assess glycolytic activity, we measured the activity of hexokinases 1 and 2 (Hk1 and Hk2), two key glycolysis enzymes that facilitate glucose entry into the glycolytic pathway. While the enzymatic activity of hexokinases and the protein levels of Hk1 and glyceraldehyde-3-phosphate dehydrogenase (Gapdh) remained largely unchanged in KO ESCs, compared to control cells (Fig. 3J, K), the sustained ATP levels (Fig. 3C) suggest a potential increase in glycolytic flux or enhanced fatty acid oxidation (FAO). This possibility is further supported by the transcriptional upregulation of metabolic genes, as presented in the following section. Collectively, our data demonstrate that the inactivation of CI function via knockout of its core subunit induces severe oxidative and reductive stress, characterized by elevated ROS levels, a markedly reduced NAD+/NADH ratio, and pronounced reduction in MMP, while overall cellular ATP levels are maintained through compensatory mechanisms.

### 4. Reductive stress caused by decreased NAD+/NADH ratio suppresses proliferation of KO ESCs

We hypothesized that the elevated ROS levels observed in KO ESCs might be responsible for their reduced proliferation. To address this possibility, we treated both KO and control ESCs with various pharmaceutical antioxidants and counted the cells 72 hours later. Treatment with optimal concentrations of the general antioxidants N-acetylcysteine (NAC, 2 mM) or Ebselen (Ebs, 5 µM), as well as the mtROS-targeting antioxidants MnTBAP (25 µM) or EUK134 (2.5 µM), failed to rescue the proliferation defect in KO ESCs (Fig. 4A-D). These treatments had no effect on cell viability, as the percentage of dead cells remained comparable to that of untreated controls. Notably, scavenging ROS with NAC or Ebs, but not with MnTBAP or EUK134, significantly reduced the proliferation of control ESCs (Fig. 5A-D), suggesting that basal cytoplasmic ROS levels are required for normal ESC proliferation. These findings indicate that the high ROS levels in KO ESCs are not direct cause of their proliferation impairment. Importantly, treatment with the SOD-mimetic MnTBAP, but not with EUK134, very mildly increased the proliferation of KO ESCs [69, 70]. This demonstrates that mitochondrial O₂^•⁻^ as well as ROS levels are not primary regulators of KO ESC proliferation.

**Fig. 4.**
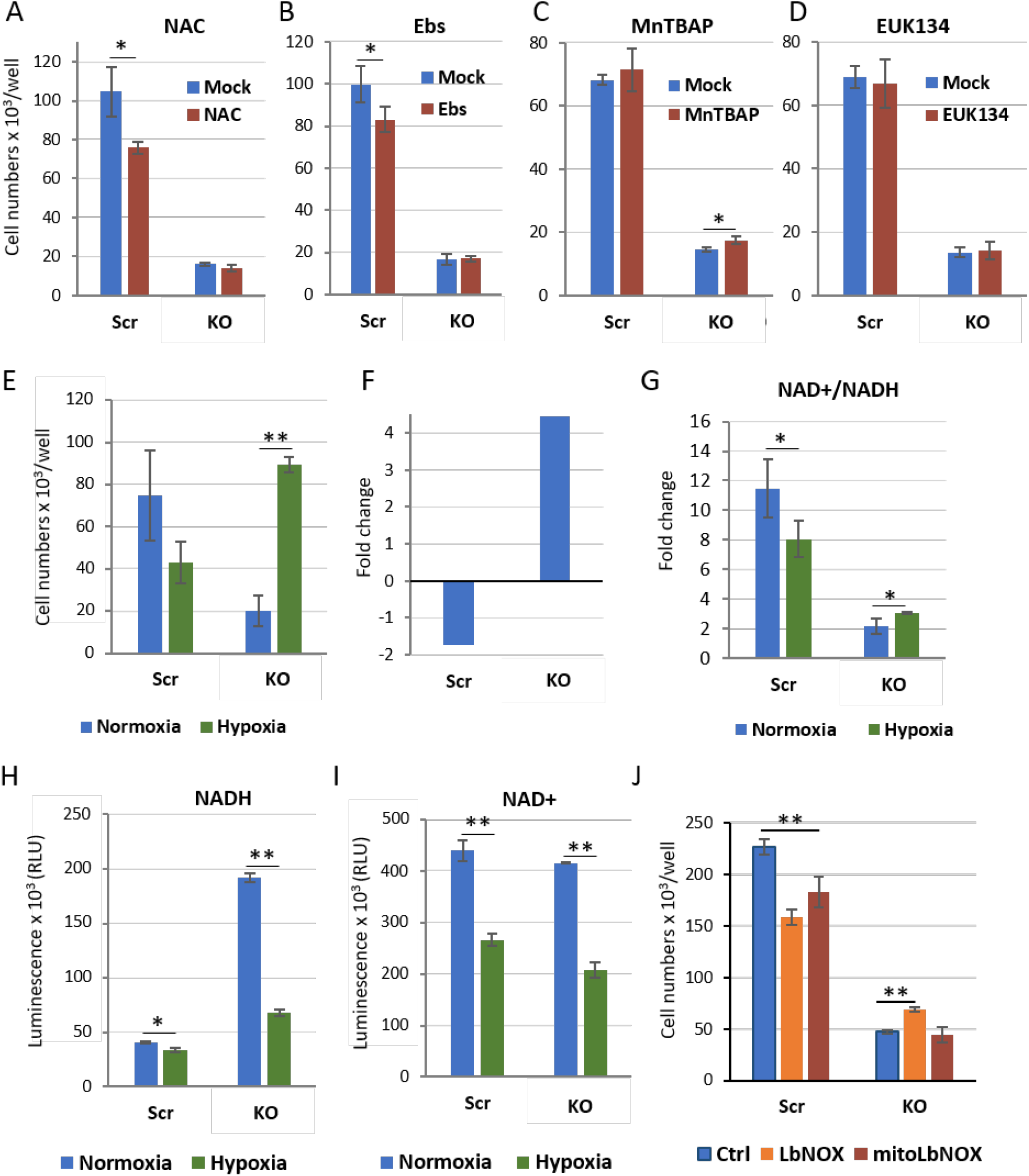
Hypoxia rescues impaired proliferation of KO ESCs by increasing the NAD+/NADH ratio. (A-D) ROS are not responsible for proliferation defects in KO ESCs. Treatment with the antioxidants (A) 2 mM N-acetyl-cysteine (NAC), (B) 5 μM Ebselen (Ebs), (C) 25 μM MnTBAP, or (D) 5 μM EUR-134 for 72 hours failed to restore proliferation in KO ESCs. NAC and Ebs treatments significantly reduced proliferation in control Scr ESCs; mean ± SD; n=3, *p<0.05. E) Hypoxia (3% O_2_, 72 hours) restores proliferation of KO ESCs but not of Scr controls; mean ± SD; n=3; **p<0.001. (F) Relative fold-changes in proliferation of KO and Scr control ESCs under hypoxia (3% O_2_, 72 hrs). G) Hypoxia (3% O_2_, 72 hours) significantly increases the NAD+/NADH ratio in KO ESCs while having the opposite effect in Scr controls; mean ± SD; n=3; *p<0.05. H) NADH levels were significantly decreased in KO ESCs and, to a much lesser extent, in Scr controls under hypoxia compared to normoxia; mean ± SD; n=3; **p<0.001. I) NAD+ levels were significantly reduced under hypoxia in both Scr and KO ESCs compared to normoxia; mean ± SD; n=3; *p<0.05, **p<0.001. J) Genetic elevation of the NAD+/NADH ratio through overexpression of LbNOX or mitoLbNOX restores proliferation of KO ESCs; mean ± SD; n=3; **p<0.001. LbNOX-mediated increase of the NAD+/NADH ratio also significantly enhanced proliferation of control Scr ESCs.

**Fig. 5.**
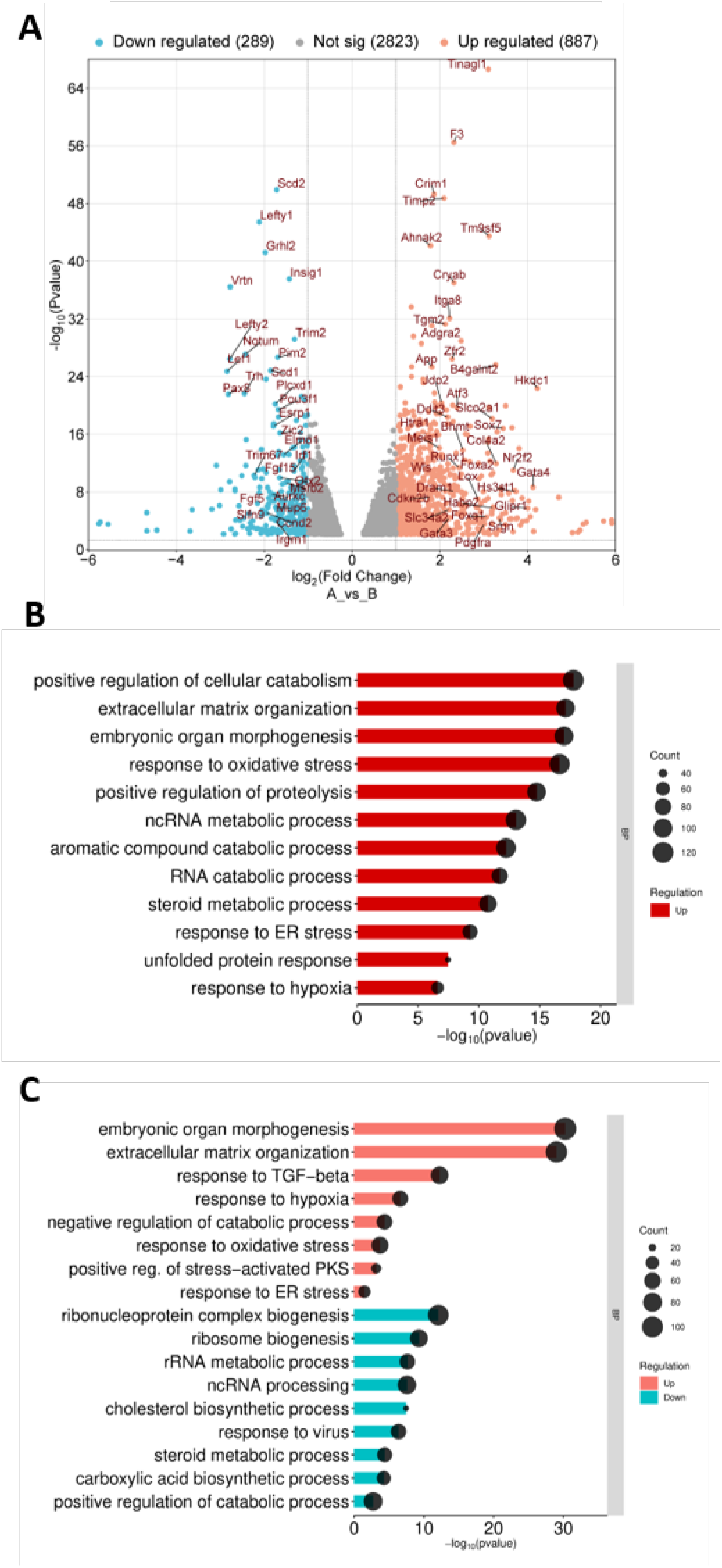
Transcriptome analysis identifies widespread gene expression changes in KO ESCs, featuring upregulation of stress response and morphogenesis pathways alongside downregulation of ribonucleoprotein biogenesis and steroid biosynthesis. (A) Volcano plot of differential gene expression analysis identifies 289 downregulated and 887 upregulated genes in KO versus control ESCs. (B) Significantly enriched gene ontology (GO) terms in KO ESCs identified by over-representation analysis (ORA). (C) Top significantly upregulated and downregulated biological processes (BP) in KO ESCs compared to Scr controls, identified by gene ontology enrichment analysis (GOEA).

An important regulator of pluripotent stem cell proliferation is hypoxia (1-5% O_2_), which activates hypoxia inducible factor-1 (HIF-1) and activates the transcription of glycolytic genes essential for cell growth [71, 72]. We found that culturing cells under hypoxic conditions (3% O_2_) for 72 h significantly increased the proliferation of KO ESCs, while control ESCs proliferated nearly twofold slower compared to normoxia (Fig. 4E, F). Specifically, KO ESCs proliferated more than four times faster under hypoxia than under normoxic conditions (Fig. 4F). These results indicate that of HIF-1 activation promotes the expression of genes involved in the positive regulation of proliferation, which are suppressed in KO ESCs under normoxic conditions.

Next, we sought to determine whether reductive stress, characterized by a decreased NAD+/NADH ratio, could be the primary signal responsible for the suppressed proliferation of KO ESCs. To determine whether NADH levels are altered under hypoxic conditions, we measured them in control and KO ESCs cultured under 3% O₂ condition. Importantly, hypoxia significantly increased the NAD+/NADH ratio in KO ESCs (Fig. 4G), primarily due to a substantial 64% decrease in NADH levels, compared to a modest 17% reduction in controls (Fig. 4H). It is noteworthy that the level of NAD+ decreased proportionally in both Scr (39%) and KO (49%) ESCs under hypoxia (Fig. 4I). Thus, the reduction in NADH concentration and the ensuing increase in the NAD+/NADH ratio correlate with the restoration of proliferation in KO ESCs under hypoxia. These data indicate that cellular NADH, rather than NAD+ depletion, regulate the proliferation of ESCs with impaired CI function. It is also important to note that sufficient NAD+ levels are maintained in both control and KO ESCs to sustain viability, suggesting that this oxidized metabolite, crucial for anabolic processes, is steadily supplied via compensatory dehydrogenase systems, such as the glycerol-3-phosphate or malate-aspartate shuttles, even when the dehydrogenase function of CI is inhibited.

To obtain mechanistic evidence that elevated NADH levels are responsible for the suppressed proliferation of KO ESCs, we evaluated the effect of directly oxidizing NADH using the bacterial H_2_O-forming NADH oxidase from *Lactobacillus brevis*, LbNOX [51]. Ectopic expression of cytoplasmic LbNOX, but not mitochondria-targeted (mitoLbNOX) partially (by 30%) and significantly rescued the impaired proliferation of KO ESCs (Fig. 4J). Surprisingly, mitoLbNOX failed to rescue proliferation, suggesting that mitochondrial NADH accumulation is not involved in the signaling pathway regulating proliferation in this context. Instead, an elevated cytosolic NADH pool plays a critical role in restricting KO ESC proliferation. Thus, our data suggest that reductive stress, mediated by elevated NADH levels and a consequent drop in the cytosolic NAD+/NADH ratio, acts as a signal suppressing proliferation in Ndufs1-deficient ESCs.

### 5. *Ndufs1* knockout upregulates catabolic genes and suppresses biosynthetic pathways

To elucidate the molecular consequences of *Ndufs1* inactivation in ESCs, we performed RNA-seq analysis, comparing the transcriptomes of KO and control Scr ESCs. Quality control confirmed high read quality and high inter-sample correlation (Fig. S3D). Differential gene expression analysis revealed 289 downregulated and 887 upregulated genes in KO ESCs (Fig. 5A, Supplementary Dataset 1). Over-representation analysis (ORA) highlighted several significantly enriched gene ontology (GO) terms, including positive regulation of catabolic processes, steroid and RNA catabolism, embryonic organ morphogenesis, extracellular matrix organization, and stress responses (Fig. 5B, Supplementary Dataset 2, Supplementary Table 1). Gene set enrichment analysis (GSEA) further revealed the upregulation of gene sets associated with embryonic organ morphogenesis, extracellular matrix organization, responses to oxidative stress and hypoxia, as well as negative regulation of catabolic processes (Fig. 5C and Fig. S3E-G, Supplementary Table 2). Conversely, genes involved in ribonucleoprotein complex biogenesis, as well as in RNA, cholesterol, and carboxylic acid biosynthesis, were significantly downregulated (Fig. 5C and Fig. S3H, Supplementary Table 2). Notably, gene sets related to oxidative, chemical, integrated stress (ISR), and unfolded protein (UPR) responses were significantly upregulated upon Ndufs1 function loss in ESCs (Fig. 7A, Fig. 8A, Supplementary Table 2). Interestingly, among the ETC complexes, only genes encoding subunits of CIV were significantly transcriptionally up- and downregulated in KO ESCs (Supplementary Dataset 1), indicating a lack of global compensatory upregulation of other ETC complex components.

The RNA-seq data further indicated that while most of glycolytic genes remained unchanged, glucose transporters (*Slc2a1/Glut1*, *Slc2a3/Glut3*, *Slc2a4/Glut4*) and hexokinases (*Hk1*, *Hk2*) were transcriptionally upregulated (Fig. S4A, B, D), suggesting a possible compensatory increase in glycolytic flux in KO ESCs. Furthermore, compensatory ATP production may also arise from enhanced mitochondrial long-chain fatty acid oxidation (FAO), as evidenced by the pronounced upregulation of most elements of this pathway (Fig. S4C). This catabolic shift can supply FADH_2_ to CII, potentially bypassing the CI deficiency. Based on the transcriptomic profiles, we propose that RNA and cholesterol catabolism are activated upon CI deficiency, contributing to a general suppression of anabolic processes, including RNA and protein synthesis.

Interestingly, the RNA-seq analysis revealed a 3- to 4-fold transcriptional upregulation of genes associated with extraembryonic (XEN) endoderm differentiation, such as *Foxa2*, *Gata4*, *Gata6*, *Sox17*, and *Sox7*, as well as of genes associated with epithelial-mesenchymal transition (EMT) (Fig. 6A). Quantitative RT-PCR confirmed the upregulation of *Foxa2*, *Sox17*, and *Gata6* transcripts (Fig. 6B). However, immunoblotting showed no detectable increase in Sox7 or Gata4 protein levels, and Foxa2 protein remained undetectable in both control and KO ESCs (Fig. 6C). These data suggest that while CI inactivation triggers transcription of early endodermal genes, the concomitant energy and metabolite deficiency, alongside activated catabolic pathways and the ISR pathway, may lead to targeted RNA degradation and impaired translation, preventing the synthesis of non-essential proteins for this particular cell state.

**Figure 6.**
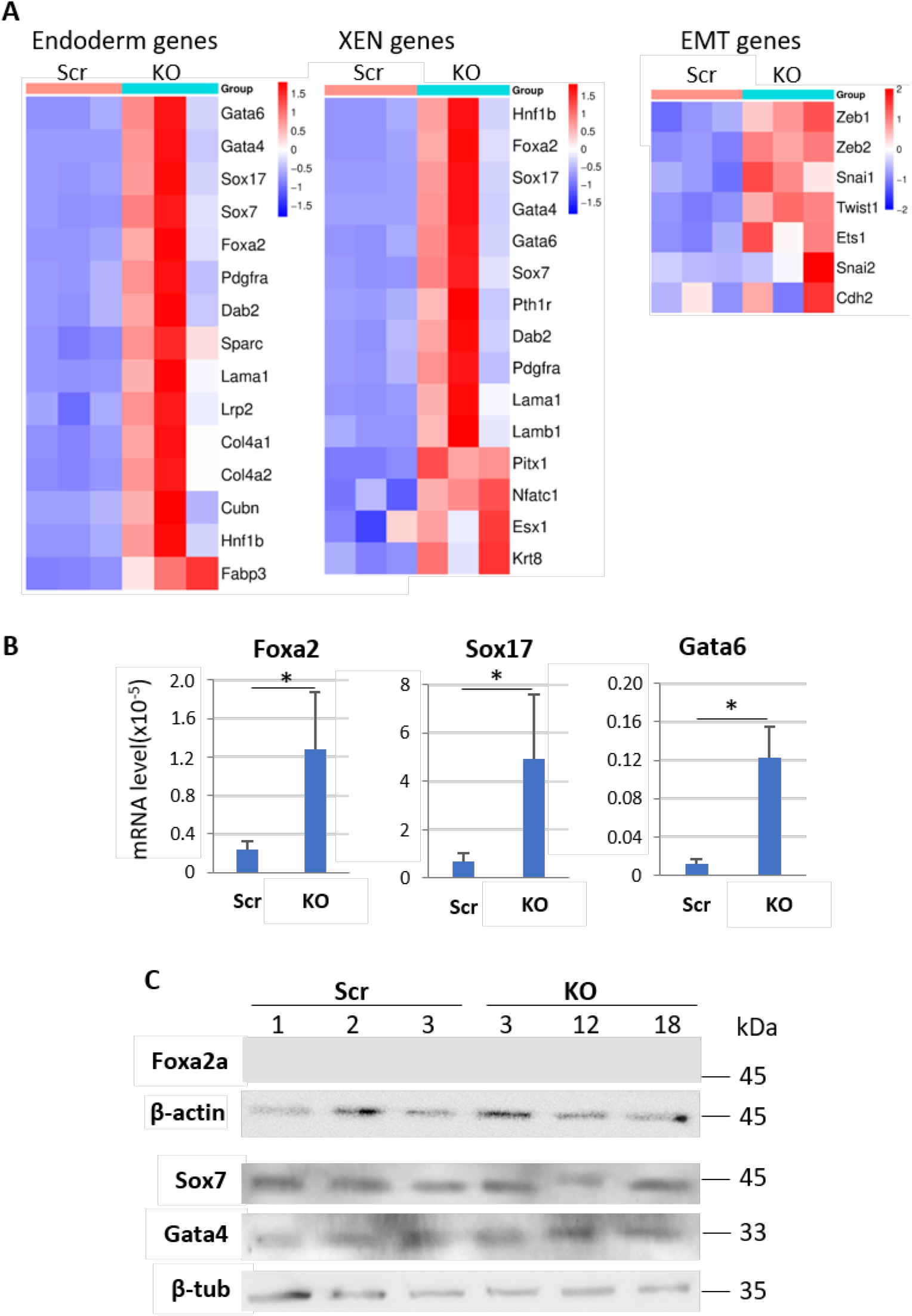
RNA-seq analysis identifies significant upregulation of transcripts associated with endoderm differentiation and epithelial-mesenchymal transition (EMT) in KO ESCs. (A) Heatmaps of DEGs show upregulation of endoderm, extraembryonic endoderm (XEN), EMT markers. (B) qRT-PCR analysis confirms significant transcriptional upregulation of the endodermal markers *Foxa2, Sox17,* and *Gata6* in KO ESCs; mean ± SD; n=3, *p <0.001. (C) Western blot analysis reveals the absence of Foxa2 protein expression and unchanged protein levels of Sox7 and Gata4 in KO ESCs compared to Scr controls. β-Actin or β-tubulin (β-tub) were used as loading controls.

### 6. Ndufs1 inactivation induces the ISR and GSH-related antioxidant genes while suppressing NRF2/SOD2 axis in ESCs

The RNA-seq analysis revealed in KO ESCs a significant upregulation of genes associated with oxidative stress, UPR and ISR, corroborating the observed oxidative (elevated ROS) and reductive (increased NADH/NAD+ ratio) stresses resulting from CI inactivation. Interestingly, only some genes involved in glutathione (GSH)-mediated antioxidant response were upregulated including glutathione peroxidases (Gpx1 and Gpx3) and NAD(P)H quinone dehydrogenase 1 (Nqo1) (Fig. 7A, B). However, the protein levels of Gpx1, catalase (Cat), and glutathione synthetase (Gss) expressions remained unaltered (Fig. 7C). Notably, several enzymes responsible for generating the cellular NADPH pool, contributing to the reduced GSH production, were transcriptionally upregulated, including NADP+-dependent isocitrate dehydrogenase 2 (Idh2), malic enzyme 1 (Me1), methylenetetrahydrofolate dehydrogenase-2 (Mthfd2), and Nqo1 (Fig. 7A, B). Immunoblotting confirmed a significant increase in Idh2 protein levels, whereas lactate dehydrogenase-B (Ldhb) levels remained unchanged (Fig. 7C), suggesting a specific enhancement of Idh2 activity upon CI inactivation. These results suggest that in ESCs, CI dysfunction triggers a Gpx-centered antioxidant response to maintain hydrogen peroxide homeostasis (Fig. 7D).

**Figure 7.**
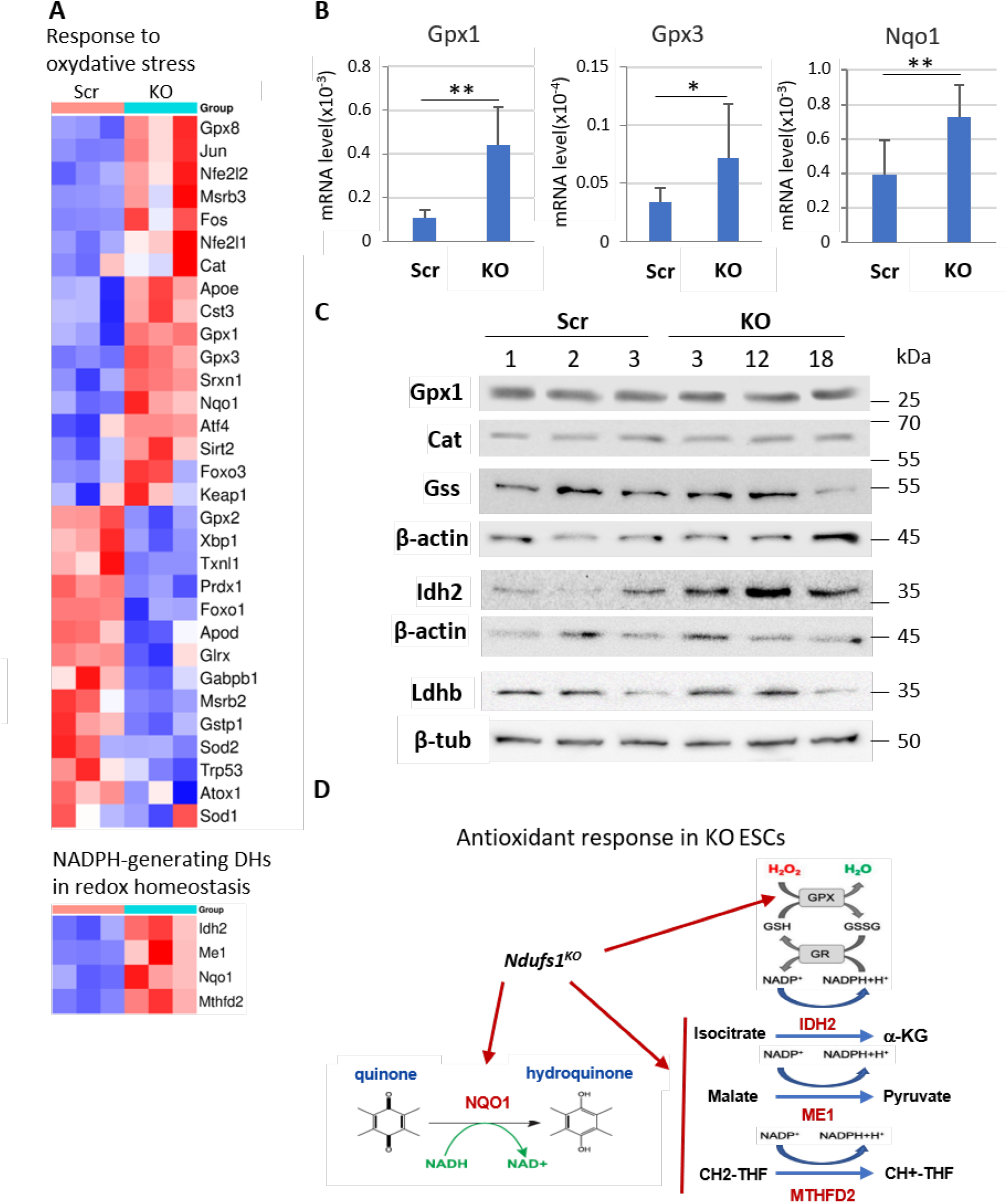
Transcriptional activation of the GPx/GSH-mediated antioxidant response in KO ESCs. (A) Heatmaps of DEGs show upregulation of antioxidant genes *Gpx1, Gpx3, Gpx8, Cat, and Nqo1*, alongside downregulation of *Gpx2* and *Sod2*. (B) qRT-PCR analysis confirms the significant upregulation of *Gpx1, Gpx3*, and *Nqo1* transcripts in KO ESCs. mean ± SD; n=3 *p <0.05, **p <0.01. (C) Western blot analysis shows unchanged levels of Gpx1, catalase (Cat), and glutathione synthetase (Gss), and Ldhb enzymes, while elevated Idh2 protein levels in KO ESCs. β-Actin or β-tubulin (β-tub) were used as loading controls. (D) Schematic model of the antioxidant response in KO ESCs.

Surprisingly, we observed that elevated mtROS levels resulting from CI dysfunction are not perceived as detrimental by ESCs but instead they may play a physiological role in supporting self-renewal and viability. Specifically, RNA-seq and Western blot data revealed that mitochondrial Sod2 levels are significantly downregulated in KO ESCs (Figs. 7A and 8). Furthermore, the Keap1/Nrf2 pathway, a master antioxidant system regulating Sod2, was suppressed in KO ESCs. This was evidenced by significant downregulation of nuclear respiratory factor-2 (Nrf2/Nfe2l2) and its phosphorylated form (ph-Nrf2), alongside an upregulation of Keap1 protein (Fig. 8). Accordingly, peroxisome proliferator-activated receptor gamma coactivator 1-α (Pgc-1α), a key regulator of mitochondrial biogenesis and Sod2 expression, together with nuclear respiratory factor 1 (Nrf1) [73–75] was also significantly downregulated at the protein level (Fig. 8). Thus, even under elevated ROS conditions, Ndufs1-dependent CI inactivation suppresses the Pgc-1α and Keap1/Nrf2 axes. This suggests that maintaining elevated mitochondrial O₂^•⁻^ levels, potentially through reduced Sod2 expression, might be an important requirement for ESCs to sustain mitochondrial signaling/function under these conditions, though the functional significance of this observation requires further investigation.

**Fig. 8.**
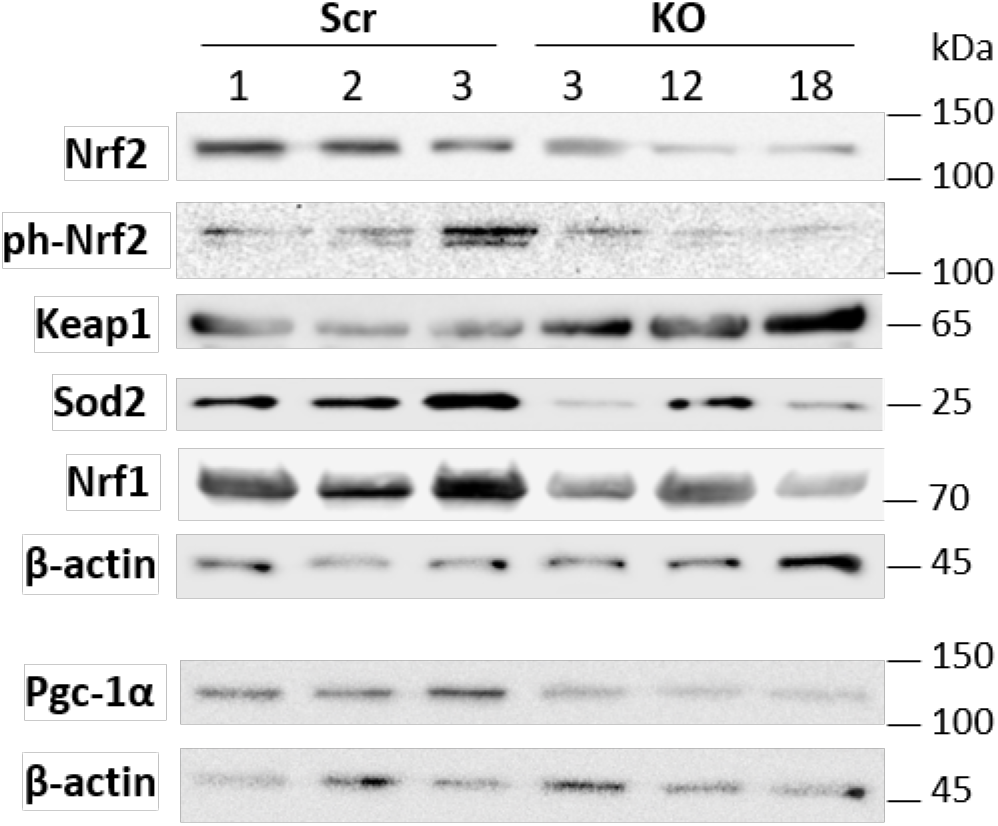
Suppressed mitochondrial antioxidant Nrf2/Keap1/Sod2 stress response pathway in KO ESCs. Western blot analysis shows significantly reduced levels of Nrf2, phospho-Nrf2 (ph-Nrf2), Nrf1, Pgc-1α, and Sod2 alongside increased Keap1 in KO ESCs. β-Actin was used as a loading control.

On the other hand, *Ndufs1* inactivation leads to upregulation of key ISR factors, such as Atf4, Atf3, Ddit3/CHOP, Ddit4/REDD1, Cebpb/C/EBPβ, Ppp1r15b, Asns, Herpud1, Slc7a11, and Chac1 (Fig. 9A), as well as of endoplasmic reticulum (ER) stress response factors – Ern1, Atf6, Atf3, Atf4, and Ero1a (Supplementary Dataset 1). Activation of the ISR was confirmed by a robust increase of Atf4 protein levels in KO ESCs (Fig. 9B). We propose that the coupled oxidative and reductive stress in mitochondria activates the Atf4-dependent stress response pathway, as evidenced by the induction of its target genes (Fig. 9A). The ISR orchestrates cellular adaptation to various stress conditions by transiently suppressing global translation while selectively activating stress-adaptive genes. Recent studies have shown that reductive stress, induced by a low NAD+/NADH ratio, can trigger ISR to suppress global protein synthesis and induce the expression of amino acid transporters (Slc7a11, Slc7a5, Slc7a1), asparagine synthase Asns and mitochondrial metabolic-stress enzyme Hkdc1 [76–78]. Furthermore, ISR/Atf4 pathway activation may contribute to the suppression of proliferation via the upregulation of Gadd45a/b [79]. Consistent with the observed proliferation defect, multiple pro-proliferative factors (e.g. Ccnd1/2, Ccne1/2, Cdk2, Cdk6, Plk1, Cdc25a Myc, Mycl, and Mycn) were downregulated, while the anti-proliferative tumor suppressors (Cdkn1a/p21, Cdkn2a/p16, and Trp53) were upregulated upon Ndufs1 loss (Fig. S2B, Supplementary Dataset 1). Collectively, the activation of the ISR/Atf4 pathway, coupled with reduced Nrf2 and Pgc-1α activities, correlates with suppressed proliferation of ESCs.

**Fig. 9.**
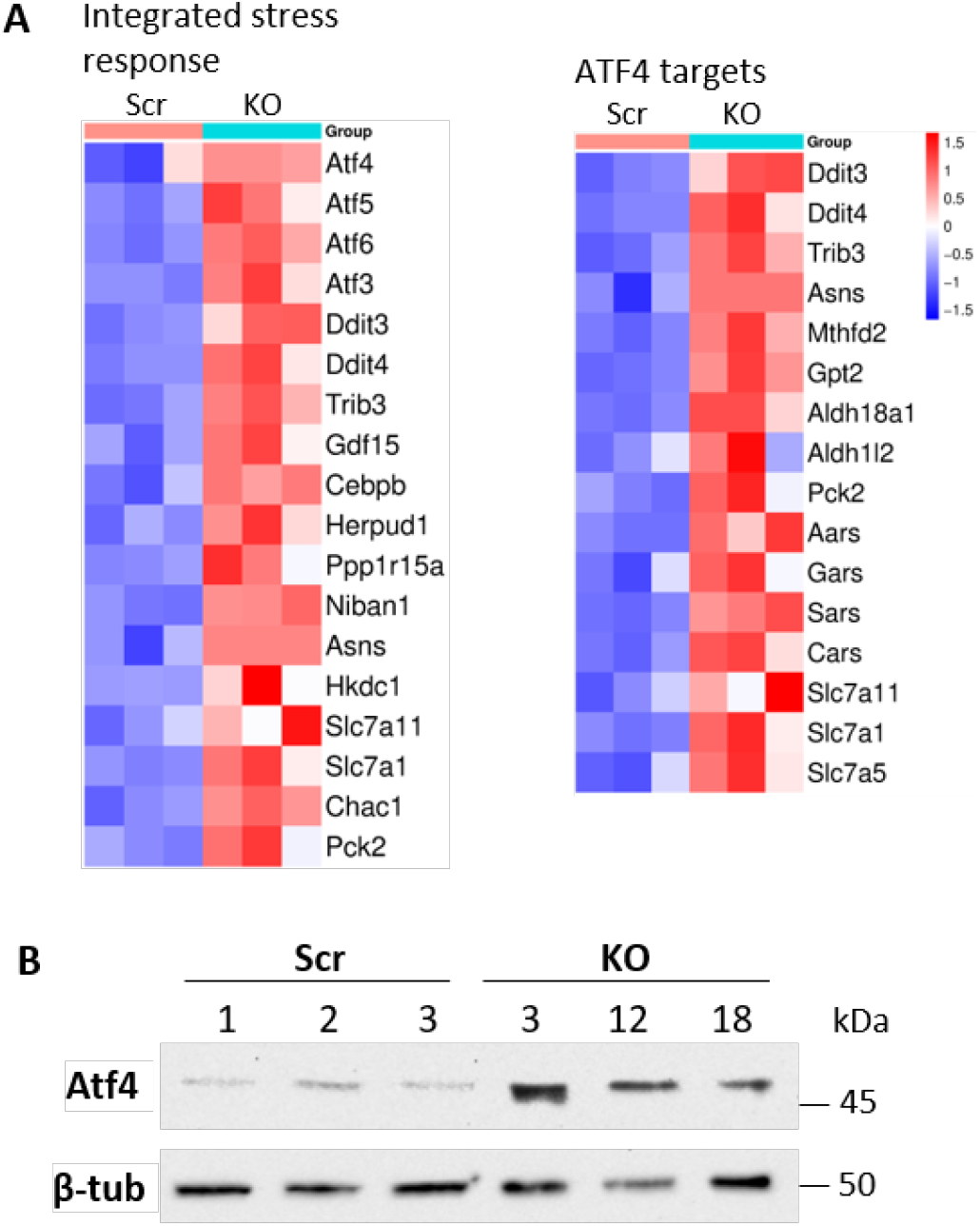
Upregulated Atf4-mediated ISR genes in KO ESCs. (A) Heatmaps of DEGs show upregulation of IRS and ATF4 target genes. (B) Markedly increased Atf4 protein levels in KO ESCs versus Scr controls. β-Tubulin (β-tub) was used as a loading control.

### 7. Hypoxia restores the Nrf2/Sod2 axis and suppresses the ISR/Atf4 response in *Ndufs1*-deficient ESCs

As described above, Nrf2/Sod2 activity is strongly suppressed, whereas ISR/Atf4 stress signaling is induced in KO ESCs. Given that hypoxia significantly improves the proliferation of KO ESCs, we assessed the level of these proteins under hypoxic conditions (3% O_2_). Importantly, under hypoxia, Nrf2 levels were restored, even slightly upregulated, while Pgc1-α and Sod2 levels in KO ESCs reached levels comparable to those in Scr controls (Fig. 10A). In contrast, Nrf1 protein levels remained similarly reduced in KO ESCs under both normoxic and hypoxic conditions. This suggests that the stabilization of Nrf2/Pgc1-α/Sod2 axis is a specific adaptive response to hypoxia in the KO ESCs context, likely contributing to the restoration of proliferation. These finding further imply that activation of Sod2 may play a pro-proliferative role under NADH-induced reductive stress, through an unknown mechanism, which requires further investigation.

**Fig. 10.**
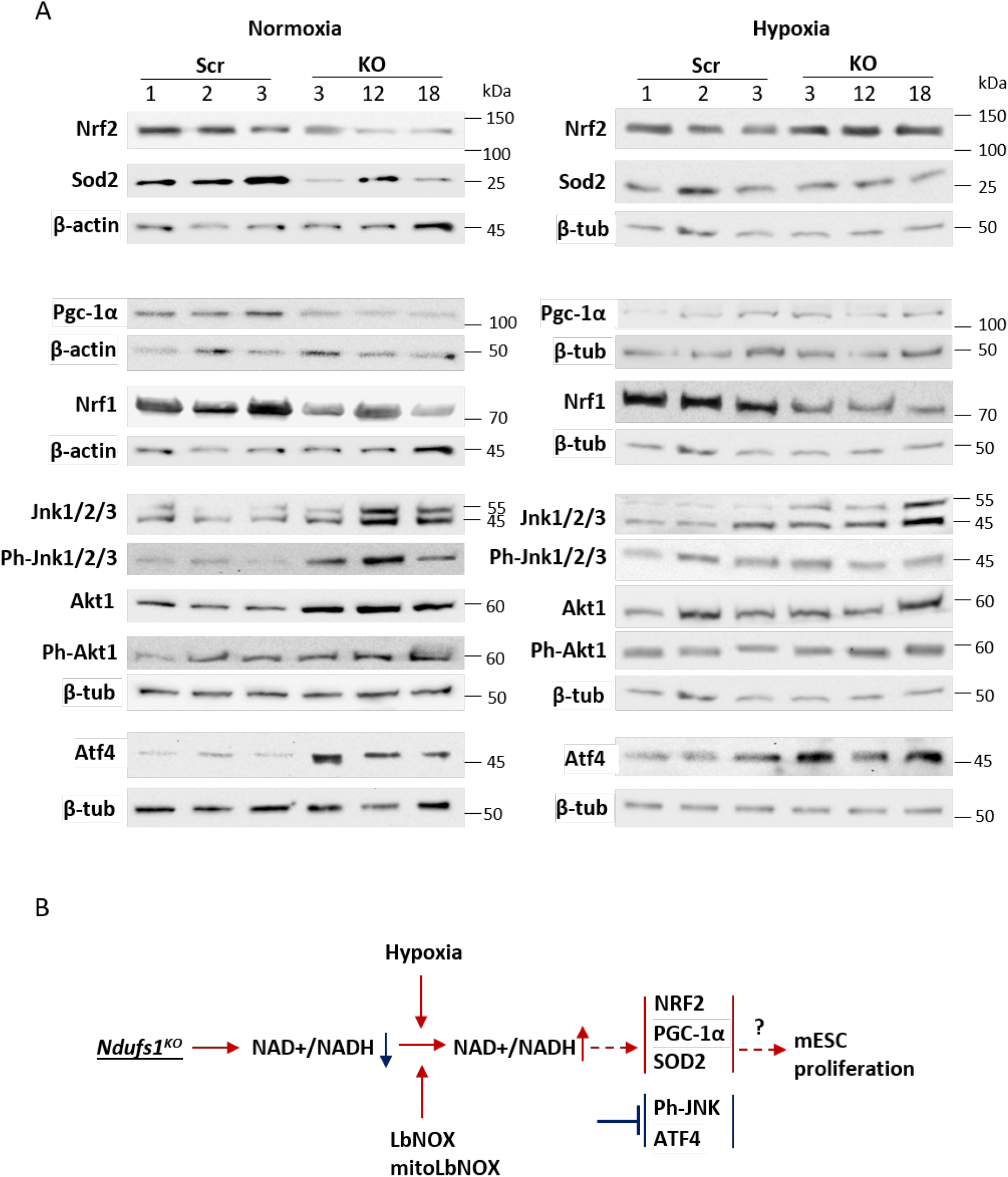
Hypoxia suppresses reductive stress-induced ISR and restores Nrf2 and Sod2 levels in KO ESCs. (A) Western blot analysis shows significantly reduced Atf4 protein levels in KO ESCs under hypoxia compared to normoxia, relative to Scr controls. Slightly elevated Nrf2 levels, unaltered Nrf1 levels, and restored Sod2 expression were observed in KO ESCs under hypoxia compared to normoxia, relative to Scr controls. β-Actin (β-act) or β-tubulin (β-tub) were used as loading controls. (B) Proposed schematic model of the molecular mechanism regulating ESC proliferation under reductive stress caused by Ndufs1deficiency-mediated inactivation of CI of the ETC.

On the other hand, we observed that the activation of Jnk (ph-Jnk), as well as the induction of ISR/Atf4 in KO ESCs, was significantly downregulated under hypoxia compared to normoxia (Fig. 10A, Fig. S1A). Conversely, as expected, pro-proliferative Akt1/phosphor-Akt1 signaling remained unchanged between hypoxia and normoxia conditions (Fig. 10A). These results indicate that the activation of Jnk and ISR/Atf4 pathways correlates with the suppressed proliferation of ESCs, potentially acting as negative regulators of the proliferation. However, we found that treatment with ISRIB, a chemical inhibitor of ISR by reversing the effects of eIF2α phosphorylation, failed to rescue the proliferation of KO ESCs and instead further suppressed the growth of both Scr and KO ESCs (Fig. S5B, B’). This suggests that while the ISR pathway is activated, it may not be direct or sole driver of growth inhibition in this context, or that basal ISR activity is sufficient for ESC proliferation. In conclusion, we propose a model wherein reductive stress, characterized by an elevated NADH/NAD+ ratio in Ndufs1-deficient ESCs, triggers Jnk and ISR activation while suppressing the Nrf2/Pgc1-α/Sod2 axis, collectively resulting in the observed suppression of ESCs proliferation (Fig. 10B)

## Discussion

The importance of metabolic regulation in various developmental processes, including pluripotent stem cell biology, is now well established. OXPHOS represents a central bioenergetic pathway that supports essential cellular functions. Impaired ETC function underlies several debilitating mitochondrial diseases in humans, such as Leigh syndrome and MELAS. Mitochondria actively communicate with the nucleus via retrograde signaling to regulate gene expression, thereby orchestrating diverse cellular processes, including apoptosis, proliferation, cell fate specification, and cell reprogramming [80–82]. This mitochondrial–nuclear communication has also been implicated in developmental dormancy; specifically, during embryonic diapause, the epiblast undergoes OXPHOS suppression and metabolic rewiring [14, 15]. We hypothesized that at certain developmental stages, cells, including ESCs, might downregulate ETC activity in response to signals from the microenvironment or the nucleus. In this context, suppressed ETC activity could trigger specific retrograde signals that support stage-specific cellular states, including the dormant state observed during embryonic diapause. Furthermore, local oxygen tension and the consequent modulation of OXPHOS activity have been shown to significantly impact ESC properties, particularly proliferation and differentiation [83–85].

Our findings demonstrate that the genetic knockout of the Ndufs1 subunit is expected to impair CI function, leading to significant metabolic reprogramming in naïve mouse ESCs. This metabolic disruption was confirmed by a marked decrease in mitochondrial membrane potential, a decreased NAD⁺/NADH ratio, and elevated mitochondrial ROS. Interestingly, despite the predicted ETC impairment, we observed a modest rise in total ATP levels, likely attributable to compensatory pathways. However, direct measurements revealed no increase in glycolytic flux, indicating that cellular energy demands in CI-deficient ESCs are met by sustaining basal glycolysis in conjunction with presumably CII-dependent OXPHOS and potentially fatty acid oxidation (FAO). Furthermore, the deficiency led to reduced mitochondrial mass, an effect potentially mediated by the downregulation of the master mitochondrial biogenesis regulators Pgc-1α and Nrf2. The observed decrease in the NAD⁺/NADH ratio reflects an NADH-replete reductive state, a metabolic signature that promotes catabolic metabolism [86, 87]. Consistent with this, our RNA-seq data indicated a broad upregulation of genes involved in catabolic processes in KO ESCs.

Ndufs1 deficiency robustly suppressed the proliferation of mouse ESCs without altering cell cycle dynamics or increasing cell death in standard serum-based conditions, which reflect the naïve state of PSCs. Importantly, as previously shown, ground-state mouse PSCs in 2iL medium exhibit the cell-cycle alterations that reflect a diapause-like state [55]. This allowed us to hypothesize that the downregulation of Ndufs1 and overall CI activity is involved in the regulation of embryonic diapause. Despite this growth slowdown, the knockout cells maintained pluripotency across passages, as evidenced by the stable expression of pluripotency markers and the retention of differentiation potential into all three germ layers within teratomas, as well as into cardiomyocytes and primed-state EpiSCs *in vitro*. This phenotype suggests that CI inactivation induces a dormant state in naïve ESCs, thereby uncoupling proliferation from pluripotent identity. At the same time, CI-deficient teratomas exhibited an abnormal architecture characterized by thin tissue layers and large cavities. While this observation requires further investigation, it may indicate functional deficits in differentiated tissues due to cell-autonomous metabolic defects or non-cell-autonomous signaling. Supporting this notion, OXPHOS inhibition is known to disrupt cardiomyocyte differentiation [88], while the exit from pluripotency involves critical metabolic shifts [89]. Moreover, CI deficiency can trigger non-cell-autonomous mitohormetic signaling that affects systemic processes [42, 59].

Our data further support the paradigm that mitochondrial function drives proliferation, while glycolysis supports pluripotency [90]. Accordingly, *Ndufs1* knockout ESCs exhibited suppressed proliferation but maintained pluripotency, as the transcripts/proteins of most pluripotency factors remained unaltered. Notably, Klf4 protein was upregulated, while c-Myc and other members of the Myc family were downregulated. This suppression of c-Myc aligns with the observed growth arrest and stable glycolytic flux, given that c-Myc serves as a key driver of cellular anabolism and proliferation [91]. Indeed, dual deletion of c- and N-Myc in mouse ESCs attenuates anabolic programs without altering pluripotency, thereby inducing a diapause-like arrest [15, 92–94]. Conversely, high MYC levels sustain glycolysis and self-renewal in human naïve PSCs, while its loss coincides with decreased glycolysis during endoderm and mesoderm differentiation [95]. In our *Ndfus1* mutants, increased expression of the Myc repressor Foxo3 likely underpins Myc downregulation and the subsequent dormant phenotype [96, 97]. Klf4 upregulation may reinforce the pluripotency network [98] or participate in inhibiting proliferation - a mechanistic link that warrants future study.

Furthermore, our findings align with previous work showing that pharmacological disruption of OXPHOS via mitochondrial uncouplers increases ROS and attenuates ESC proliferation without compromising pluripotency or increasing cell death [99]. This underscores that in standard culture, PSCs rely on active OXPHOS to meet the bioenergetic and biosynthetic demands of rapid proliferation – a dependency shared with many cancer cell types [90]. Collectively, these insights support a model wherein during blastocyst development, mitochondrial retrograde signaling may physiologically modulate epiblast self-renewal in response to metabolic cues.

### Increased ROS does not mediate the proliferation defect in CI-deficient ESCs

To identify the specific signal suppressing the proliferation of CI-deficient ESCs, we examined altered metabolic parameters. We first ruled out ROS, given that antioxidant treatment failed to restore proliferation in KO cells and actually reduced it in the controls, indicating that basal ROS levels are required to promote PSC proliferation. Although ESCs operate under low redox stress [100] and possess high expression of antioxidant proteins [101], and despite evidence that increased ROS can drive differentiation [102–104] or inhibit Oct4 function via oxidation [105], the ROS increase in our mutants appears insufficient to explain the phenotype. Consequently, this proliferation defect likely involves alternative metabolic signals.

Ndufs1 inactivation triggers a switch in antioxidant defense, suppressing the Keap1/Nrf2 pathway while activating a glutathione peroxidase-based system reliant on mitochondrial Idh2. Idh2 promotes generation of NADPH, a key reductant for antioxidant defense [106–109]. Its upregulation in KO cells points to a compensatory NADPH/NADP⁺-dependent redox mechanism dedicated to scavenging H₂O₂ and preventing hydroxyl radical formation [110]. In this system, Gpx enzymes eliminate peroxides by oxidizing GSH to GSSG, and are then recycled by glutathione reductase using NADPH. Together, these findings highlight the specific activation of an NADPH-dependent Gpx1/Gpx3 pathway involving Idh2, Me1, and Mthfd2 to counteract oxidative stress in CI-deficient ESCs (Fig. 7D).

Importantly, Idh2 activity is integral to ETC biogenesis and function. Its loss impairs mitochondrial respiration, reducing the expression of CI-III and elevating ROS in diverse models [111, 112]. Specifically, Idh2 facilitates the assembly of CI and other OXPHOS complexes [111]. Thus, its upregulation in *Ndufs1* mutants may represent a compensatory mechanism to stabilize the ETC architecture, a possibility that warrants further investigation. Notably, Idh2 is itself a transcriptional target of the core pluripotency factors Oct4, Sox2, and Nanog [12], further reinforcing the metabolic-pluripotency nexus.

It was surprising that in *Ndufs1*-mutant ESCs, there was a robust downregulation of Sod2 at both the mRNA and protein levels. The concurrent downregulation of Sod2 and suppression of the Nrf2 pathway are particularly notable, as Nrf2 is a primary transcriptional regulator of *Sod2* [113, 114]. This suggests that CI-derived O₂^•⁻^ actively suppresses the Nrf2/Sod2 axis in this context. Giving that Sod2 is the sole mitochondrial enzyme converting O₂^•⁻^ to H₂O₂, its reduction suggests a metabolic shift where mitochondria sustain high O₂^•⁻^ but low H₂O₂ levels. In this scenario, O₂^•⁻^ conversion to H₂O₂ likely occurs in the cytosol, either spontaneously or via Sod1, with subsequent H₂O₂ clearance by the cytosolic Gpx system. Unlike O₂^•⁻^, H₂O₂ is a key signaling molecule; consequently, limiting its mitochondrial production via decreased Sod2 expression may protect respiratory complexes and suppress oxidative stress-related signaling events that are known to destabilize OXPHOS complexes [111]. Furthermore, a decrease in Sod2 expression/function has been associated with the dissolution of mitochondrial supercomplexes, which in turn reduces respiration [115, 116], potentially forming a protective feedback loop that regulates the stability of ETC complexes. We therefore hypothesize that CI-deficient ESCs adapt to elevated mitochondrial O₂^•⁻^ by inhibiting the Nrf2-Sod2 pathway to maintain a low-H₂O₂ environment, thereby protecting mitochondrial integrity and supporting cell viability. The precise trigger for this downregulation of the Nrf2-Sod2 axis in the context of CI deficiency, however, remains to be elucidated.

### Hypoxia improves the proliferation of CI-deficient ESCs by elevating the NAD⁺/NADH ratio

We found that hypoxia significantly improves the proliferation rate of Ndufs1-deficient ESCs. This aligns with the established role of Hif1-α in promoting pluripotency and self-renewal by inhibiting spontaneous differentiation [83, 84, 117–120]. Hif-1α and Hif-2α upregulate a network of genes involved in proliferation (c-Myc, Ccnd1), survival (Vegf), pluripotency (Oct4, Nanog), antioxidant defense (Nrf2, ATM), and glycolysis (Glut1, Pgk, Ldha, Adrp, Adm) [121–128]. While hypoxia-induced glycolytic activation could contribute to this proliferative rescue, our data argue against ATP as the limiting factor: total ATP is already elevated in KO cells, and further glycolytic stimulation with nicotinic acid (NA) fails to improve proliferation (Fig. S5C). Instead, the hypoxia-mediated increase in the NAD⁺/NADH ratio may be the critical metabolic signal restoring proliferation.

This observation implies that hypoxia rescues the proliferation of Ndufs1-deficient ESCs suggests that an elevated NAD⁺/NADH ratio under low oxygen conditions is a critical signal. This implies that the high NADH levels accumulated due to impaired CI activity directly inhibit proliferation. NADH, generated by glycolysis, the TCA cycle, and FAO, serves as central electron donor for OXPHOS and biosynthesis [129, 130]. Notably, the presence of excess aspartate in the culture medium failed to rescue this proliferation defect (Fig. S5D), indicating that the growth arrest in CI-deficient ESCs is primarily driven by NADH-mediated metabolic toxicity rather than a shortage of biosynthetic precursors. In cancer cells, mitochondrial NADH buildup feedback-inhibits TCA cycle enzymes, thereby slowing growth, though serine-derived NADH production via Mthfd2/Mthfd2L escapes this regulation [131, 132]. Consequently, pathological NADH accumulation can suppress key pathways governing in cell growth [131–133]. We found that in KO ESCs, *Mthfd2* is upregulated, which may provide high steady-state levels of mitochondrial NADH. The growth-inhibitory role of NADH is further supported by the observation that ectopic expression of LbNOX partially rescues the proliferation defect, while PDH activation - which promotes NADH production - further suppresses the proliferation of Ndufs1-deficient ESCs (Fig. S5E).

Finally, we observed that while the ISR is activated in *Ndufs1*-deficient ESCs, it is not directly involved in growth inhibition, even though its activity was significantly reduced under hypoxic conditions. The ISR is a highly conserved mechanism that orchestrates the cellular response to various extreme conditions, modulating protein synthesis and selectively promoting the expression of pro-survival genes [76, 77, 134]. Its outcome is not uniform and varies depending on the nature, duration, and intensity of the stressor. In *Ndufs1*-deficient ESCs, the ISR may trigger a reduction in the translation of non-essential genes, explaining the absence of protein expression for early endodermal genes, despite their transcriptional activation. We demonstrate that the activation of the ISR/Atf4 pathway, coupled with the suppression of the Nrf2/Pgc-1α/Sod2 axis, correlates with impaired proliferation in CI-deficient cells (Fig. 10B). This coordinated response appears to protect pluripotent cell identity and modulate proliferation through metabolic and redox rewiring triggered by reductive stress.

Collectively, our study defines a critical role for CI activity in the proliferation of naïve ESCs, which is separable from their differentiation potential. We establish that CI deficiency induces a metabolic state characterized by NADH accumulation, suppression of the Nrf2/Sod2 antioxidant axis, and activation of a compensatory Idh2/Gpx system. This state is not driven by an ATP deficit or oxidative stress but is directly linked to a high NADH/NAD^+^ ratio. Hypoxia or enforced LbNOX expression, by reversing this ratio, restores proliferation, further exposing NADH as a central retrograde signal. We propose a model wherein mitochondrial CI activity, through its impact on the NAD⁺/NADH balance, functions as a metabolic rheostat for ESC proliferation, potentially mirroring a physiological checkpoint during early development.

## List of abbreviations

CI: Complex I
DHE: dihydroethidium
EpiLCs: epiblast-like stem cells
EpiSCs: epiblast stem cells
Ebs: Ebselen
ESCs: embryonic stem cells
ETC: electron transport chain
FAO: fatty acid oxidation
GSEA: gene set enrichment analysis
GSH: glutathione
gRNAs: guide RNAs
ISR: integrated stress responses
IMM: inner mitochondrial membrane
KO: knockout
LbNOX: NADH oxidase from *Lactobacillus brevis*
LIF: leukemia inhibitory factor
MMP: mitochondrial membrane potential
MSR: MitoSOX Red
MTR: Mitotracker Red
mtDNA: mitochondrial DNA
NAC: N-acetylcysteine
Ndufs1: NADH:ubiquinone oxidoreductase core subunit S1/ core 75 kDa subunit
ORA: over-representation analysis
OXPHOS: oxidative phosphorylation
qRT-PCR: quantitative real-time reverse transcription polymerase chain reaction
ROS: reactive oxygen species
SL: serum/LIF
Scr: scrambled
UPR: unfolded protein response

## Declarations

## Ethics approval and consent to participate

All animal procedures were conducted in strict accordance with the animal protection legislation of the Russian Federation and were approved as a humane use of laboratory animals by the Institutional Ethical Committee of the Institute of Cytology RAS (Project Title is « Study of the role of Ndufs1 in maintaining pluripotency of mouse embryonic stem cells», Approval No. 07/25, approval date: Sept. 16, 2025). All animal experiments were conducted and reported in strict compliance with the ARRIVE guidelines 2.0.

## Consent for publication

Not applicable.

## Availability of data and materials

All data generated or analyzed during this study are included in this published article and its Supplementary Information files. The RNA sequencing data generated during this study have been deposited in the NCBI Gene Expression Omnibus (GEO) repository and are accessible under the accession number GSE345877. Other raw datasets are available from the corresponding author on reasonable request.

## Competing interests

The authors declare no competing interests.

## Funding

This research was funded by the Russian Science Foundation grant № 26-14-00126.

## Author Contributions

T.Y.S. developed and conducted the majority of experiments, analyzed the data, created the artwork, and edited the manuscript; A.A.K. performed experiments, including the development of gene knockout ESCs and primary phenotypic analysis; I.B.N., E.V.S., S.V.P., V.D. performed experiments; N.D.A. performed flow cytometry analyses; A.N.T. provided resource support and edited the manuscript; S.A.S. conceived the study, developed and managed the project, provided grant support, created the artwork, wrote the original draft, and edited the manuscript. All authors have read and agreed to the published version of the manuscript.

## Acknowledgments

We thank Dr. Alla N. Shatrova for assistance with flow cytometry. The authors declare that they have not used AI-generated work in this manuscript.

## Notes

https://www.ncbi.nlm.nih.gov/geo/query/acc.cgi?acc=GSE345877

### Competing Interest Statement

The authors have declared no competing interest.

### Summary of Updates

Dear bioRxiv Team, Author names and grant information information were corrected. Please correct the author names and grant information on the title page https://www.biorxiv.org/content/10.64898/2026.10.26.761997v1. Author names should appear without the suffixes Sr. of Jr. This research was funded by the Russian Science Foundation, grant no. 26-14-00126, https://rscf.ru/project/26-14-00126 Kind regards, Sergey A. Sinenko

