## Supplementary material for "Complex I is essential for embryonic stem cell proliferation through NADH-mediated redox signaling": Supplimentary Figures

### Figure S1

A

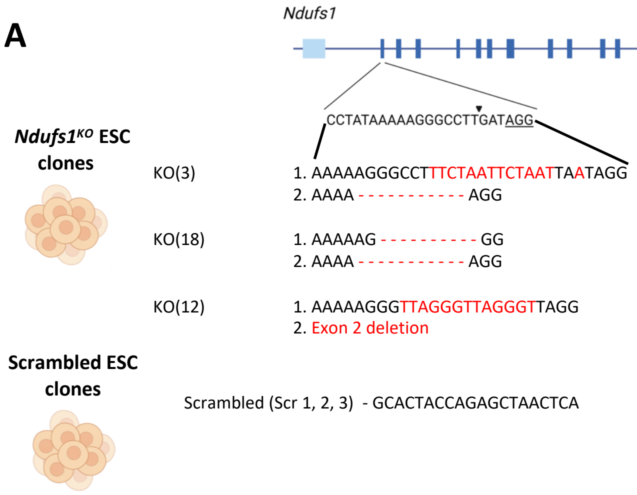


B

Chromosome counts in *Ndufs1*$^{KO}$ lines:

| | KO (3) | KO (12) | KO (18) | Scr (1) |
|---|---|---|---|---|
| % correct Chr. Content N=10 | 80% | 80% | 90% | 80% |

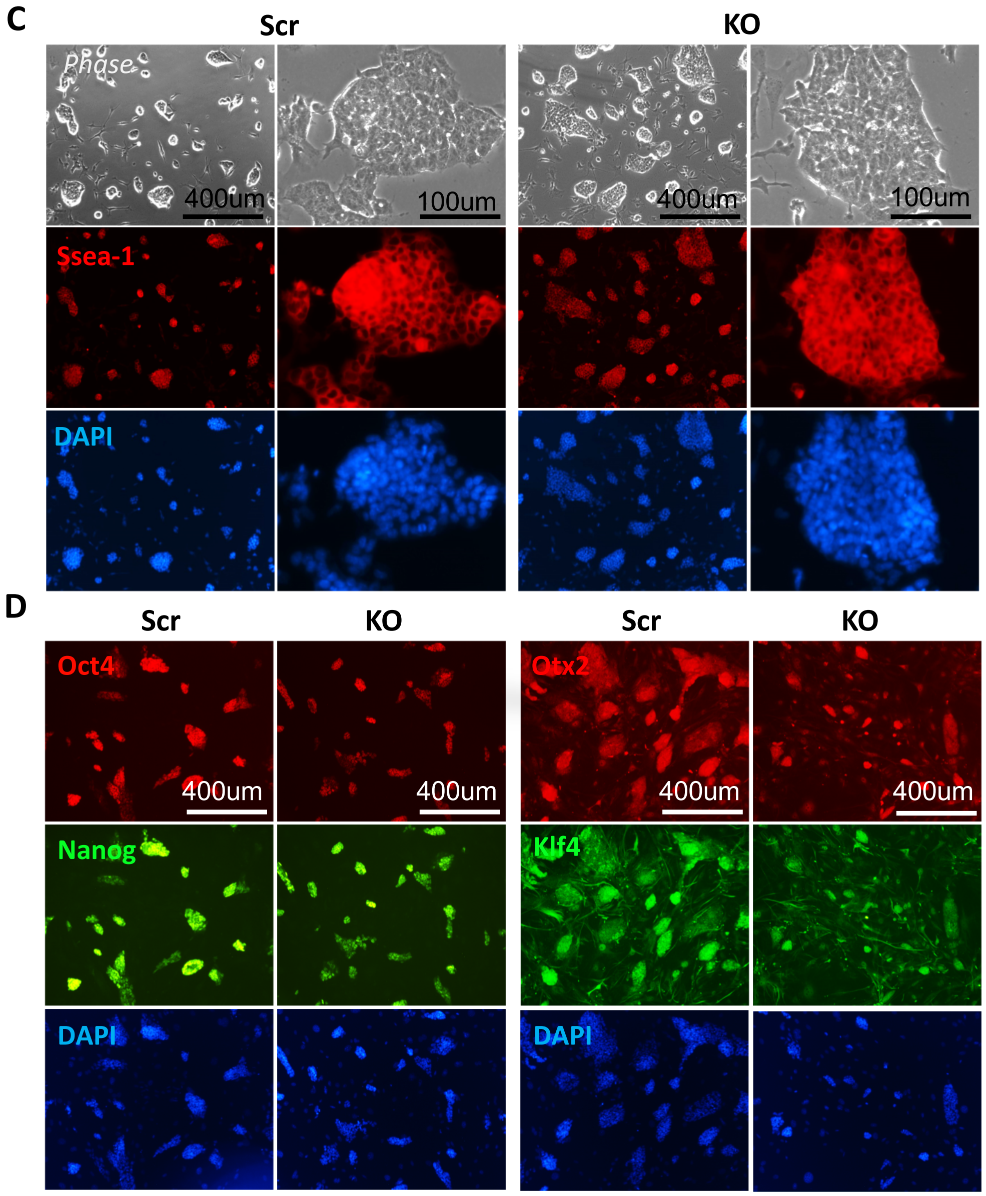

**Figure S1. (A,B) *Ndufs1$^{KO}$* (KO) alleles. (A)** Schematic representation of *Ndufs1* gene locus and sequences of *Ndufs1$^{KO}$* (KO) alleles (3, 18, 12) and control scrambled guide RNA oligonucleotide (Scr 1, 2, 3). **(B)** Proportion of correct chromosome numbers in KO alleles and control scrambled Scr (1) as revealed by DAPI staining and fluorescent microscopy (N=10).

**(C,D) Pluripotency marker expression and cell morphology are unaltered in KO ESCs.**

**(C)** Phase contrast microscopy and immunostaining for the pluripotency marker SSEA-1 (red) show intact morphology of KO ESC clones compared to Scr controls.

**(D)** Immunocytochemistry analysis confirms that the expression of pluripotency markers Oct4, Nanog, and Klf4 (color-coded) is unaffected in KO ESCs relative to Scr controls. Cells were cultured on a feeder layer of mitomycin-C-treated mouse embryonic fibroblasts (MEFs) in the SL medium. Nuclei were counterstained with DAPI (blue). Scale bar: 400 µm.

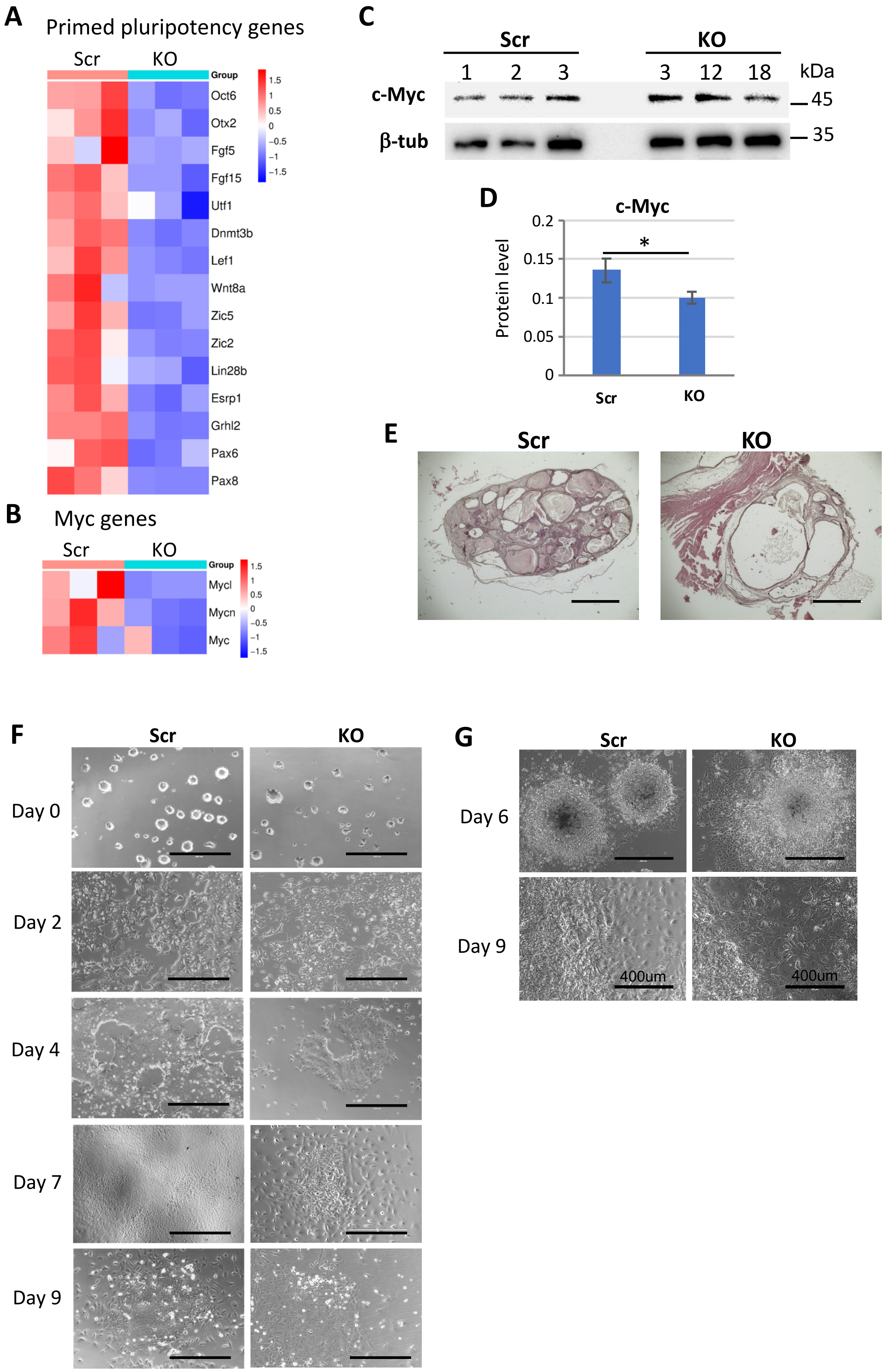
A
Primed pluripotency genes
Scr
KO
Group
1.5
1
0.5
0
−0.5
−1
−1.5
Oct6
Otx2
Fgf5
Fgf15
Utf1
Dnmt3b
Lef1
Wnt8a
Zic5
Zic2
Lin28b
Esrp1
Grhl2
Pax6
Pax8
B
Myc genes
Scr
KO
Group
Mycl
Mycn
Myc
C
Scr
KO
1
2
3
3
12
18
kDa
c-Myc
45
β-tub
35
D
c-Myc
Protein level
0.2
0.15
0.1
0.05
0
*
Scr
KO
E
Scr
KO
F
Scr
KO
Day 0
Day 2
Day 4
Day 7
Day 9
G
Scr
KO
Day 6
Day 9
400um
400um

**Figure S2. KO ESCs remains unaltered differential potential. (A)** Significantly downregulated expression of genes associated with primed pluripotency in KO ESCs. **(B)** The analysis also revealed a significant transcriptional downregulation of Mycl and Mycn genes in KO mESCs. **(C)** Western blot analysis of c-Myc expression in KO and Scr control ESC clones. α-Tubulin (Tub) was used as a loading control. **(D)** Quantification of c-Myc band intensity normalized to α-Tubulin of Western blot (panel C); mean ± SD; n=3 independent clones, $*p<0.001$. **(E)** Altered morphology of teratomas derived from KO mESCs. Histological sections of teratomas formed from control (Scr) and *Ndufs1*$^{KO}$ mESCs. Teratomas from *Ndufs1*$^{KO}$ cells exhibit an altered structure, characterized by extensive fluid-filled cavities. Scale bar: 1000 µm.

**(F)** KO ESCs differentiated into EpiSCs with efficiency comparable to Scr controls. Phase-contrast images documenting the morphological progression of Scr and KO cells from the naive pluripotent state (day 0) to the primed EpiSC state (day 9). Scale bar: 400 µm. **(G)** KO ESCs differentiated into cardiomyocytes with efficiency comparable to Scr controls. Phase-contrast micrographs documenting dynamics of Scr and KO cell differentiating from the pluripotent state (day 0) to cardiomyocytes (day 9). Scale bar: 400 µm.

Figure S3

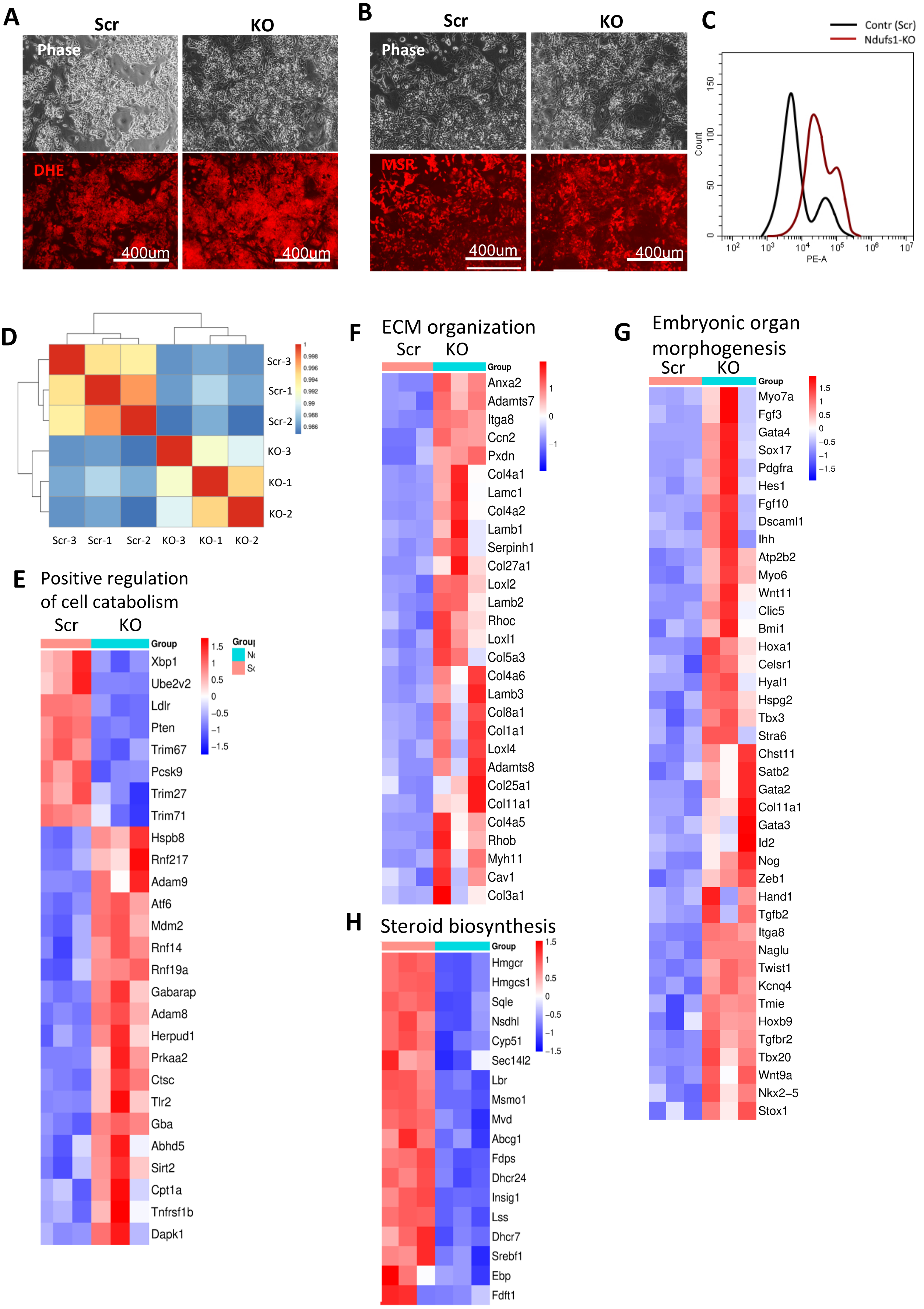
A
Scr
KO
Phase
DHE
400um
400um
B
Scr
KO
Phase
MSR
400um
400um
C
Contr (Scr)
Ndufs1-KO
Count
PE-A
D
Scr-3
Scr-1
Scr-2
KO-3
KO-1
KO-2
E
Positive regulation of cell catabolism
Scr
KO
Group
Xbp1
Ube2v2
Ldlr
Pten
Trim67
Pcsk9
Trim27
Trim71
Hspb8
Rnf217
Adam9
Atf6
Mdm2
Rnf14
Rnf19a
Gabarap
Adam8
Herpud1
Prkaa2
Ctsc
Tlr2
Gba
Abhd5
Sirt2
Cpt1a
Tnfrsf1b
Dapk1
F
ECM organization
Scr
KO
Group
Anxa2
Adamts7
Itga8
Ccn2
Pxdn
Col4a1
Lamc1
Col4a2
Lamb1
Serpinh1
Col27a1
Loxl2
Lamb2
Rhoc
Loxl1
Col5a3
Col4a6
Lamb3
Col8a1
Col1a1
Loxl4
Adamts8
Col25a1
Col11a1
Col4a5
Rhob
Myh11
Cav1
Col3a1
G
Embryonic organ morphogenesis
Scr
KO
Group
Myo7a
Fgf3
Gata4
Sox17
Pdgfra
Hes1
Fgf10
Dscaml1
Ihh
Atp2b2
Myo6
Wnt11
Clic5
Bmi1
Hoxa1
Celsr1
Hyal1
Hspg2
Tbx3
Stra6
Chst11
Satb2
Gata2
Col11a1
Gata3
Id2
Nog
Zeb1
Hand1
Tgfb2
Itga8
Naglu
Twist1
Kcnq4
Tmie
Hoxb9
Tgfbr2
Tbx20
Wnt9a
Nkx2–5
Stox1
H
Steroid biosynthesis
Group
Hmgcr
Hmgcs1
Sqle
Nsdhl
Cyp51
Sec14l2
Lbr
Msmo1
Mvd
Abcg1
Fdps
Dhcr24
Insig1
Lss
Dhcr7
Srebf1
Ebp
Fdft1

**Figure S3. Elevated $O_2^{\bullet-}$ levels in KO ESCs**. **(A)** Representative phase-contrast (phase) and fluorescence micrographs showing increased total cellular $O_2^{\bullet-}$ levels in KO ESCs compared to Scr controls, as determined by DHE staining (red). Scale bar: 400 µm. **(B)** Representative phase-contrast (phase) and fluorescence micrographs showing increased mitochondrial $O_2^{\bullet-}$ levels in KO ESCs, as determined by MSR staining (red). Scale bar: 400 µm. **(C)** Flow cytometry analysis reveals significantly elevated mitochondrial superoxide in KO ESCs. Cytograms show two cell populations with low and high MSR fluorescence.

**(D-H) Transcriptome analysis identifies widespread gene expression changes in KO ESCs. (D)** Quality control and overview of RNA-seq data. Sample-to-sample correlation. Heatmap depicting the pairwise correlation of global gene expression profiles across all samples, demonstrating high reproducibility within experimental groups. Heatmap of DEGs shows upregulation of cell catabolism **(E)**, extracellular matrix (ECM) organization **(F)**, and embryonic organ morphogenesis processes **(G)**, as well as downregulation of steroid biosynthesis **(H)** processes.

Figure S4

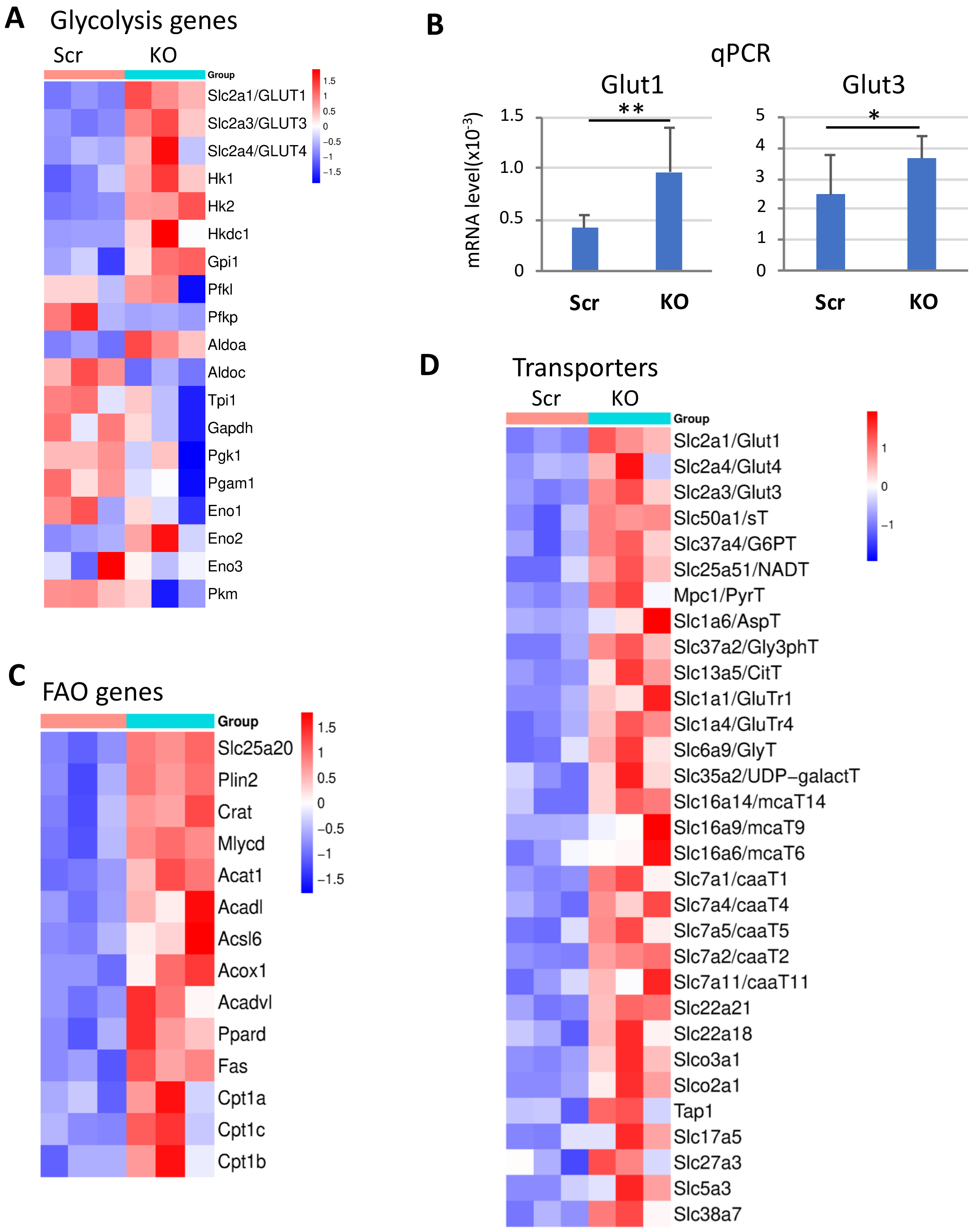


**Figure S4. Transcriptomic analysis identifies significant upregulation of fatty acid oxidation (FAO) and membrane transporter genes in KO ESCs. (A)** Heatmap of differentially expressed genes (DEGs) shows upregulation of some glycolytic genes. **(B)** qRT-PCR analysis validates the significant upregulation of glucose transporters *Glut1* (*Slc2a1*) and *Glut3* (*Slc2a3*) in KO ESCs. *p <0.05, **p <0.01. **(C)** Heatmap reveals a coordinated upregulation of FAO genes. **(D)** Heatmap displays widespread upregulation of genes encoding various membrane transporters.

Figure S5

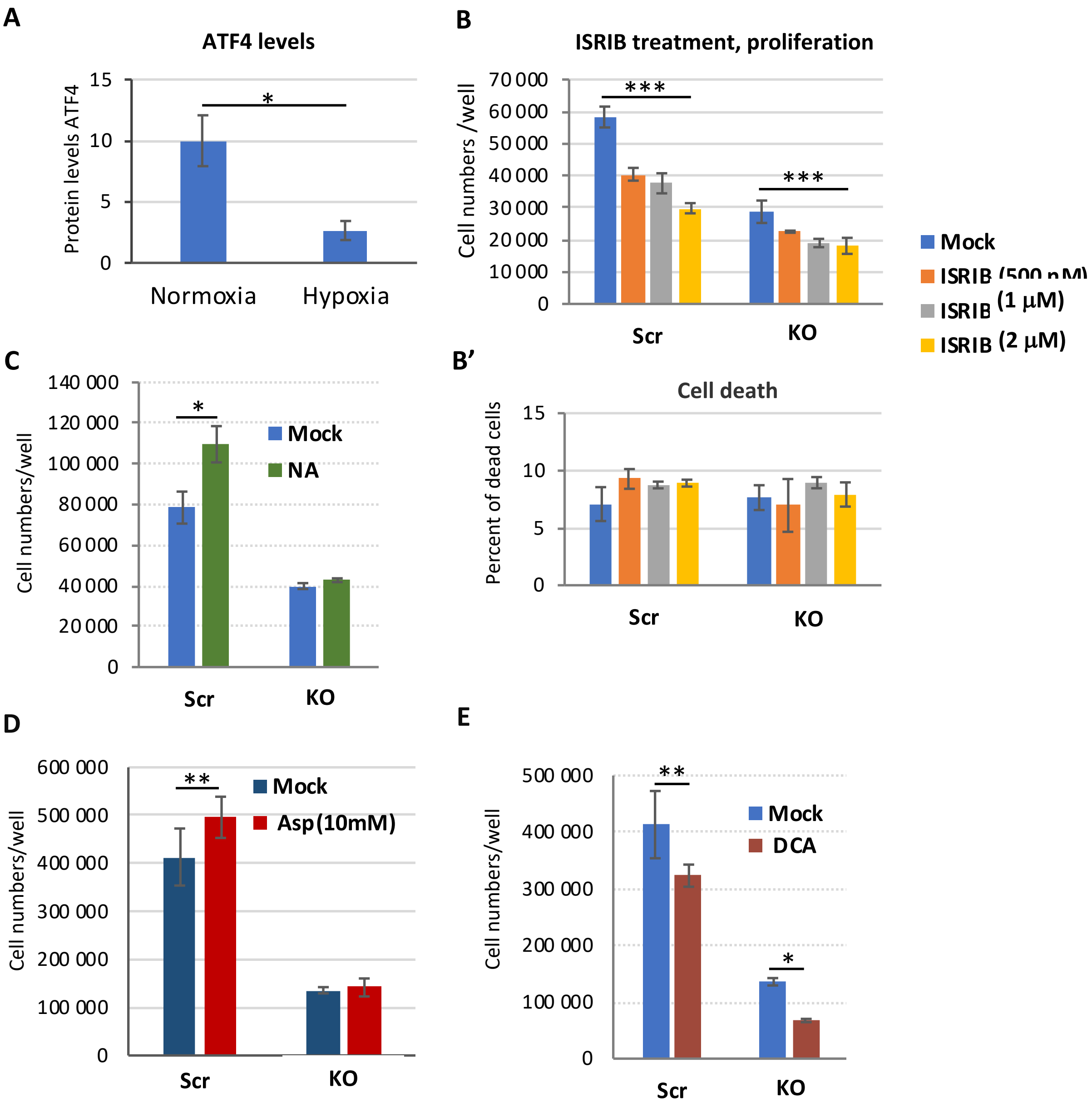


**Figure S5. The role of ISR/ATF4 signaling in the regulation of ESC proliferation. (A)** Quantification of ATF4 band intensity normalized to α-Tubulin from the Western blot experiment shown in Figure 7A. Data are presented as mean ± SD; n=3 independent clones. $p < 0.001$. **(B)** Treatment with the integrated stress response inhibitor (ISRIB) at concentrations of 0.5, 1, or 2 µM for 72 hours suppresses proliferation in both control Scr and KO ESCs. Data are presented as mean ± SD; n=3. *$p < 0.005$. **(B')** Percent of dead cells in the experiment of the panel **B**.

**(C-E) Activation of the TCA cycle, but not glycolysis, suppresses proliferation of KO ESCs. (C)** Treatment with 10 µM nicotinic acid (NA) for 72 hours increased the proliferation of Scr ESCs but failed to rescue the impaired proliferation in KO ESCs; mean ± SD; n=3; *$p<0.001$. **(B)** Treatment with 10 mM dichloroacetate (DCA) for 72 hours significantly suppressed proliferation in both KO ESCs and Scr control ESCs; mean ± SD; n=3; **$p<0.05$, *$p<0.001$. **(E)** Excess of aspartate (Asp, 10 mM) failed to rescue the impaired proliferation in KO ESCs, but slightly increases proliferation of control ESCs; mean ± SD; n=3; **$p<0.05$.
